# Rho of plant GTPases with geranylgeranylation motif modulate monoterpene indole alkaloid biosynthesis in *Catharanthus roseus*

**DOI:** 10.1101/2023.09.04.554920

**Authors:** Dikki Pedenla Bomzan, Anuj Sharma, Pamela Lemos Cruz, Ines Carqueijeiro, Léo Bellenger, Avanish Rai, Akshay Kumar Thippesh, S.C. Venkatesha, Durgesh Parihar, Eric Ducos, Vincent Courdavault, Dinesh A. Nagegowda

**Affiliations:** Molecular Plant Biology and Biotechnology Lab, CSIR-Central Institute of Medicinal and Aromatic Plants, Research Centre, Bengaluru-560065; Academy of Scientific and Innovative Research (AcSIR), Ghaziabad-201002, India; Université de Tours, EA2106 Biomolécules et Biotechnologies Végétales, Tours, France; University of Horticultural Sciences, Bagalkot, India

**Keywords:** Rho of plant, CSIL motif, PGGT-I, geranylgeranylation, localization, modulation, monoterpene indole alkaloids, Cathranthus roseus

## Abstract

Rho Of Plant (ROP) GTPases function as molecular switches that control signaling processes essential for growth, development, and defense. However, their role in specialized metabolism is poorly understood. Previously, we demonstrated that inhibition of protein geranylgeranyl transferase (PGGT-I) negatively impacts the biosynthesis of monoterpenoid indole alkaloids (MIA) in *Catharanthus roseus*, indicating the involvement of prenylated proteins in signaling. Here, we show through biochemical, molecular and *in planta* approaches that specific geranylgeranylated ROPs modulate *C. roseus* MIA biosynthesis. Among the six *C. roseus* ROP GTPases (CrROPs), only CrROP3 and CrROP5, having a C- terminal CSIL motif, were specifically prenylated by PGGT-I. Additionally, both of their transcripts showed higher expression in most parts compared to other *CrROPs*. Protein- protein interaction studies revealed that both CrROP3 and CrROP5, but not CrROP2 (lacking CSIL motif), interacted with CrPGGT-I. Further, CrROP3 and CrROP5 exhibited nuclear localization, whereas CrROP2 was localized to plasma membrane. *In planta* functional studies revealed that silencing of *CrROP3* and *CrROP5* negatively affected MIA biosynthesis, while their overexpression upregulated MIA formation. In contrast, silencing and overexpression of *CrROP2* had no effect on MIA biosynthesis. Moreover, overexpression of Δ*CrROP3* and Δ*CrROP5* mutants lacking the CSIL motif failed to enhance MIA biosynthesis. Taken together, these results implicate that CrROP3 and CrROP5 have positive regulatory role on MIA biosynthesis and thus shed light on how geranylgeranylated ROP GTPases mediate the modulation of specialized metabolism in *C. roseus*.

## INTRODUCTION

*Catharanthus roseus* commonly known as Madagascar Periwinkle, is a tropical plant of the Apocynaceae family, producing a large palette of specialized metabolites called monoterpene indole alkaloids (MIA). It is the best-characterized MIA-producing plant species and some of the MIA produced by this plant have immense therapeutic values. For instance, ajmalicine and serpentine mainly produced by the roots of *C. roseus* are used in the treatment of hypertension and cardiovascular diseases (Zhao et al., 2013), and most importantly heterodimer bisindole alkaloids vinblastine and vincristine mostly accumulated in leaves have been used as antineoplastic agents (Miettinen et al., 2014; Kellner et al., 2015). The formation of MIA in *C. roseus* leaves is a highly complex process involving more than 50 biosynthetic steps. These steps are distributed across various cell layers of the leaf and involve different pathway enzymes, transcription factors (TFs), intra-/intercellular signaling molecules, and transporters (Zhao et al., 2013) (Figure 1). The MIA biosynthesis in *C. roseus* is under the regulation of diverse TFs, which include octadecanoid-responsive Catharanthus AP-2 domain (ORCA; Menke, 1999; Van Der Fits & Memelink, 2001; Li et al., 2013; Paul et al., 2017), basic helix- loop-helix (bHLH) factor (MYC2; Zhang et al., 2011), WRKY (Suttipanta et al., 2011) and bHLH iridoid synthesis (BIS; Van Moerkercke et al., 2015; 2016). In addition, MIA biosynthesis is also shown to be affected by mitogen-activated protein kinase (MPK3; Raina et al., 2013). Further, the expression of TFs and genes involved in MIA biosynthesis is induced upon exposure to methyl jasmonate (MeJA)- a phytohormone involved in defense against necrotrophs and herbivores (Aerts et al., 1994; Zhou & Memelink, 2016). Though most of the pathway genes and various transcriptional regulators involved in MIA biosynthesis have been functionally characterized, factors involved upstream of jasmonate-mediated signaling cascade are yet to be identified.

**Figure 1:**
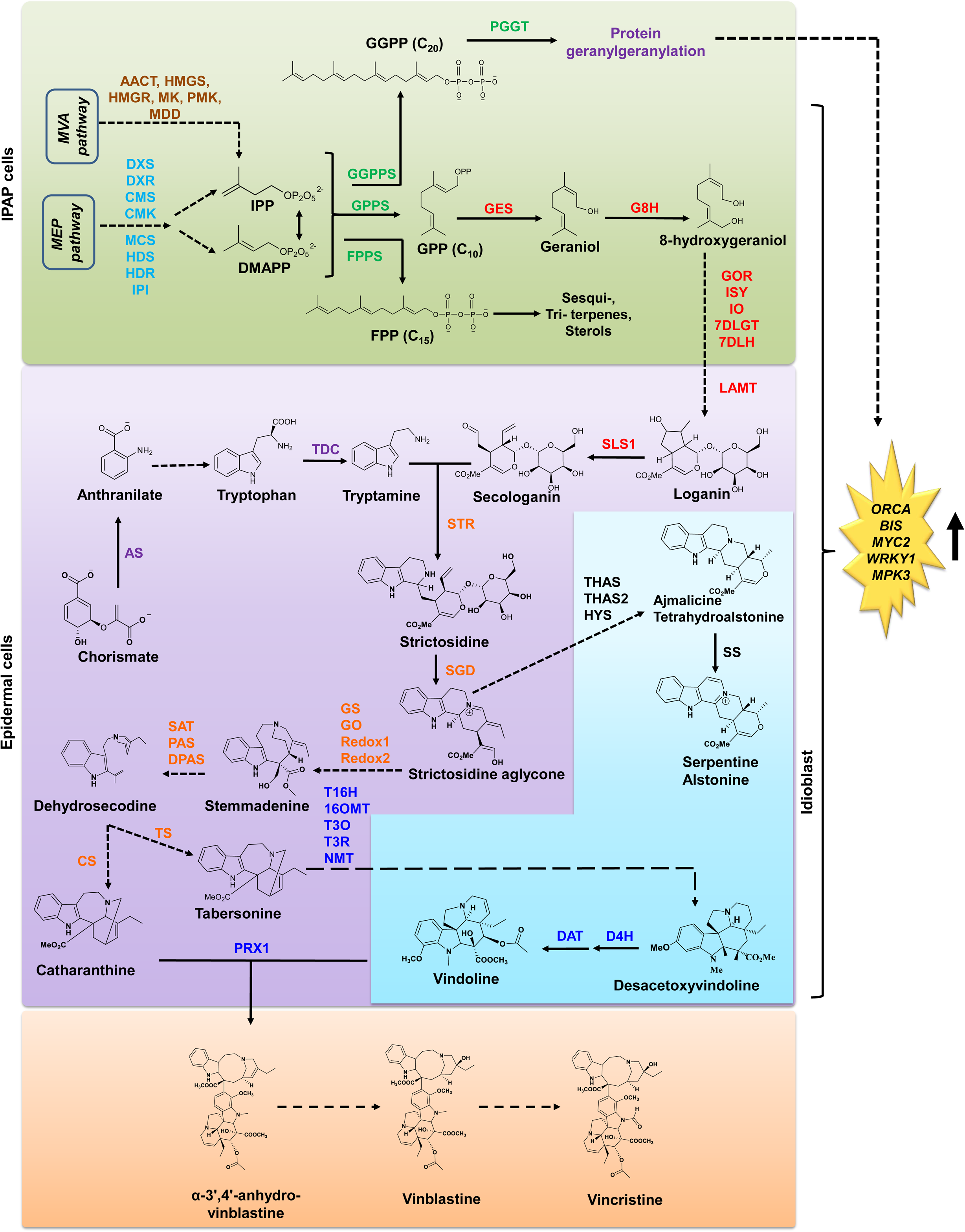
The MIA biosynthetic pathway in *C. roseus* leaves. Branch pathway enzymes are shown in different colors: MVA pathway – brown; MEP pathway - light blue; Terpenoid branch pathway – green; Iridoid pathway - red; Indole pathway – purple; Central alkaloid pathway - orange; Late alkaloid pathway - dark blue. Abbreviations: AACT, acetoacetyl-CoA thiolase; HMGS, 3-hydroxy-3-methylglutaryl-Coenzyme A synthase; HMGR, HMG-CoA reductase; MK, mevalonate kinase; PMK, phosphomevalonate kinase, MDD, mevalonate diphosphate decarboxylase; DXS, 1-deoxy-D-xylulose 5-phosphate synthase; DXR, 1-deoxy-D-xylulose5-phosphate reductoisomerase; CMS, 4-diphosphocytidyl-methylerythritol 2-phosphate synthase; CMK, 4-diphosphocytidyl- 2-C-methyl-D-erythritol kinase; MCS, 2C-methyl-Derythritol 2;4-cyclodiphosphate synthase; HDS, 1-hydroxy-2-methyl-2-(E)-butenyl-4- diphosphate synthase; HDR, 1-hydroxy-2- methyl-butenyl 4-diphosphate reductase; IPI, 3 plastid isopentenyl pyrophosphate,dimethylallyl pyrophosphate isomerase; GPPS, geranyl pyrophosphate synthase; PGGT-I; protein ger-anylgeranyl transferase I FPP, farnesyldiphosphate; FPPS, FPP synthase; GES, geraniol synthase; GGPP, geranyl-geranyl diphosphate; GGPPS, geranylgeranyl diphosphate synthase; G8H, geraniol 8-hydroxylase; GOR, 8- hydroxygeraniol oxidoreductase; ISY, iridoid synthase; IO, iridoid oxidase; 7DLGT, UDPglucose iridoid glucosyltransferase; 7DLH, 7-deoxyloganic acid 7-hydroxylase; LAMT, loganic acid methyltransferase; SLS1, secologanin synthase; TDC, tryptophan decarboxylase; STR, strictosidine synthase; AS, anthranilate synthase; SGD, strictosidine β-glucosidase; GS1, geissoschizine synthase; GO, geissoschizine oxidase; Redox1, reductive enzyme 1; Redox2, reductive enzyme 2; SAT, stemmadenine acetyl transferase; PAS, precondylocarpine acetate synthase; DPAS, dihydroprecondylocarpine acetate synthase; CS, catharanthine synthase; TS, tabersonine synthase; T16H2, tabersonine 16-hydroxylase; 16OMT, 16-hydroxytabersonine O-methyltransferase; T3O, tabersonine 3- oxidase; T3R, tabersonine 3-reductase; NMT, 16-hydroxy-2;3-dihydro-3-hydroxytabersonine N-methyltransferase; D4H, desacetoxyvindoline-4-hydroxylase; DAT, deacetylvindoline 4- Oacetyltransferase; THAS; THAS2, tetrahydroalstonine synthase; HYS, heteroyohimbine synthase; SS, serpentine synthase; PRX1, class III peroxidase 1.

It is a well-known phenomenon that when plants come in contact with elicitors, signalling cascades are activated leading to the production of secondary metabolites (Zhao et al., 2005). As a primary response to elicitors, there must be a direct and swift signaling cascade that operates before the activation of gene transcription to regulate the biosynthesis of specialized metabolites (Huchelmann et al., 2014). Post-translational modifications (PTMs) of proteins play important role in the activation of signalling cascade and have been implicated in the regulation of a number metabolic processes. PTMs of the proteins are catalyzed by the enzymatic reaction that covalently attach small chemical groups such as phosphate (phosphorylation), methyl (methylation), acetyl (acetylation), and peptides like ubiquitin (ubiquitination). In addition, protein prenylation involving farnesylation and geranylgeranylation are also important PTMs in plants. While protein farnesylation involves the activity of protein farnesyltransferase (PFT) that catalyzes the transfer of a C_15_ farnesyl moiety to the Cys residue of a protein with a C-terminal CaaX motif (C = Cys; a = aliphatic; X = Met, Gln, Ala, Cys, or Ser), protein geranylgeranylation is carried out by protein geranylgeranyltransferase (PGGT), which transfers the C_20_ geranylgeranyl lipid moiety to the target proteins (Zhang & Casey, 1996; Crowell, 2000; Crowell & Huizinga, 2009). This lipid modification plays crucial role in activation of signaling cascade required for various plant processes including abscisic acid (ABA) and auxin signaling (Johnson et al., 2005), and regulation of cell cycle (Etienne-Manneville & Hall, 2002). Besides, studies have indicated the involvement of protein geranylgeranylation in the regulation of specialized metabolism in some plants. In *Catharanthus roseus* cells, inhibition of protein geranylgeranylation negatively impacted some genes involved in the MIA pathway resulting in reduced accumulation of ajmalicine (Courdavault et al., 2005; Courdavault et al., 2009; Simkin et al., 2013). Similarly, inhibition of geranylgeranylation by the monoterpene *S*-carvone or GGti-2133 (specific inhibitor of geranylgeranylation) suppressed cellulase-induced production of sesquiterpene phytoalexin capsidiol in tobacco, implicating the involvement of geranylgeranylation in regulation of sesquiterpene biosynthesis (Huchelmann et al., 2014). In a recent study, we have demonstrated that the specific silencing and overexpression of *CrPGGT-I_*β or geranylgeranyl diphosphate synthase 2 (CrGGPPS2 that provides geranylgeranyl moiety to CrPGGT-I_β) in *C. roseus* leaves resulted in respective down- and up- regulation of MIA biosynthesis, thus establishing the importance of geranylgeranylation in MIA formation (Kumar et al., 2020). All these above studies clearly indicate that geranylgeranylation of certain, yet-to-be identified proteins might play a role in relaying downstream signals for modulation of specialized metabolism. In fact, binding proteins such as Rho-of-plants (ROPs) family of proteins are postulated to be involved in the activation of signaling resulting in the modulation of specialized metabolism (Huchelmann et al., 2014; Kurosaki & Taura, 2015).

ROP proteins are small GTPases specific to plants that belong to the Rho subfamily of the Ras superfamily, and show high similarity to the animal Rac (rat sarcoma (RAS)-related C3 botulinum toxin substrate) subfamily (Kawasaki et al., 1999; Winge et al., 2000). ROPs function as molecular switches due to changes in conformation upon GTP binding and hydrolysis (Nielsen, 2020). The conformational differences between the GTP-bound (active) and GDP-bound (inactive) states facilitate transient interactions with effector and regulatory protein (Berken & Wittinghofer, 2008; Feiguelman et al., 2018). This dynamic cycling enables ROPs to relay intracellular and extracellular stimuli in a spatially regulated manner, which results in the regulation of intracellular responses. As a result, ROPs play crucial roles in cellular and developmental processes such as pollen tube growth (which is essential for fertilization), plant cell polarity, root hair development, and auxin signaling (which regulates plant growth and development) (Gu et al., 2006; Sorek et al., 2010; Wu et al., 2011). Additionally, ROPs are also known to participate in defense responses against pathogens (Ma et al., 2017; Engelhardt et al., 2020). Although extensive research has been carried out to understand the role of ROP GTPases in different plant processes as stated above, their exact roles concerning specialized metabolism remain largely unknown. So far, there has been only a couple of reports that suggest the participation of ROPs in the modulation of plant specialized metabolism. It was shown that overexpression of *Scoparia dulcis ROP* gene (*Sdrac2)* in *Atropa belladonna*, a plant from the Solanaceae family, increased the accumulation of the specialized metabolite atropine, a tropane alkaloid (Asano et al., 2013). Similarly, overexpression of rac2 encoding a Rac GTPase in cell cultures of *Aquilaria microcarpa* activated the transcription of a sesquiterpene synthase gene encoding δ-guaiene synthase (Kurosaki & Taura, 2015). Nevertheless, these studies did not investigate the mechanism by which rac2 affects the specialized metabolism. Here, we investigated the role of ROPs in regulation of the MIA biosynthesis in the model medicinal plant *C. roseus*. Our comprehensive analysis of CrROPs through biochemical, molecular and *in planta* approaches, demonstrated that specific CrROPs (CrROP3 and CrROP5) having geranylgeranylation motif “CSIL” positively modulate MIA biosynthesis in *C. roseus*.

## RESULTS

### CrROPs and their relation to ROPs of other plants

A BLASTp search using ROPs from other plants was performed to identify ROP candidates in the *C. roseus* assembled genome (Kellner et al., 2015) (She et al., 2019). *CrROP* transcripts with very low Fragments Per Kilobase of transcript per Million mapped reads (FPKM) values were excluded and further clustering of *CrROP* sequences yielded six full-length coding sequences similar to annotated plant ROPs. All *CrROP* genes have complex gene structure with multiple introns and showed uneven distribution on four of the eight chromosomes (Figure S1). Phylogenetic analysis of CrROPs with other characterized ROPs revealed that they belong to four distinct clades (Figure 2A). CrROP3 (UHU71015) and CrROP5 (UHU71017) belong to clade-1, and exhibit highest similarity to *Scoparia dulcis* Sdrac2 (ACM07419) and *Arabidopsis thaliana* AtROP6 (Q38912), respectively (Figure 2A; Table S1). CrROP4 (UHU71016) and CrROP6 (UHU71018) belong to clade-2, and display highest relatedness to AtROP7 (AED95321) and AtROP8 (AEC10456), respectively. Whereas CrROP2 (UHU71014) and CrROP1 (UHU71013) respectively belong to clade-3 and clade-4, and show highest similarity to AtROP9 (NP_194624) and Am-rac1 (AEQ62558) (Figure 2A).

**Figure 2.**
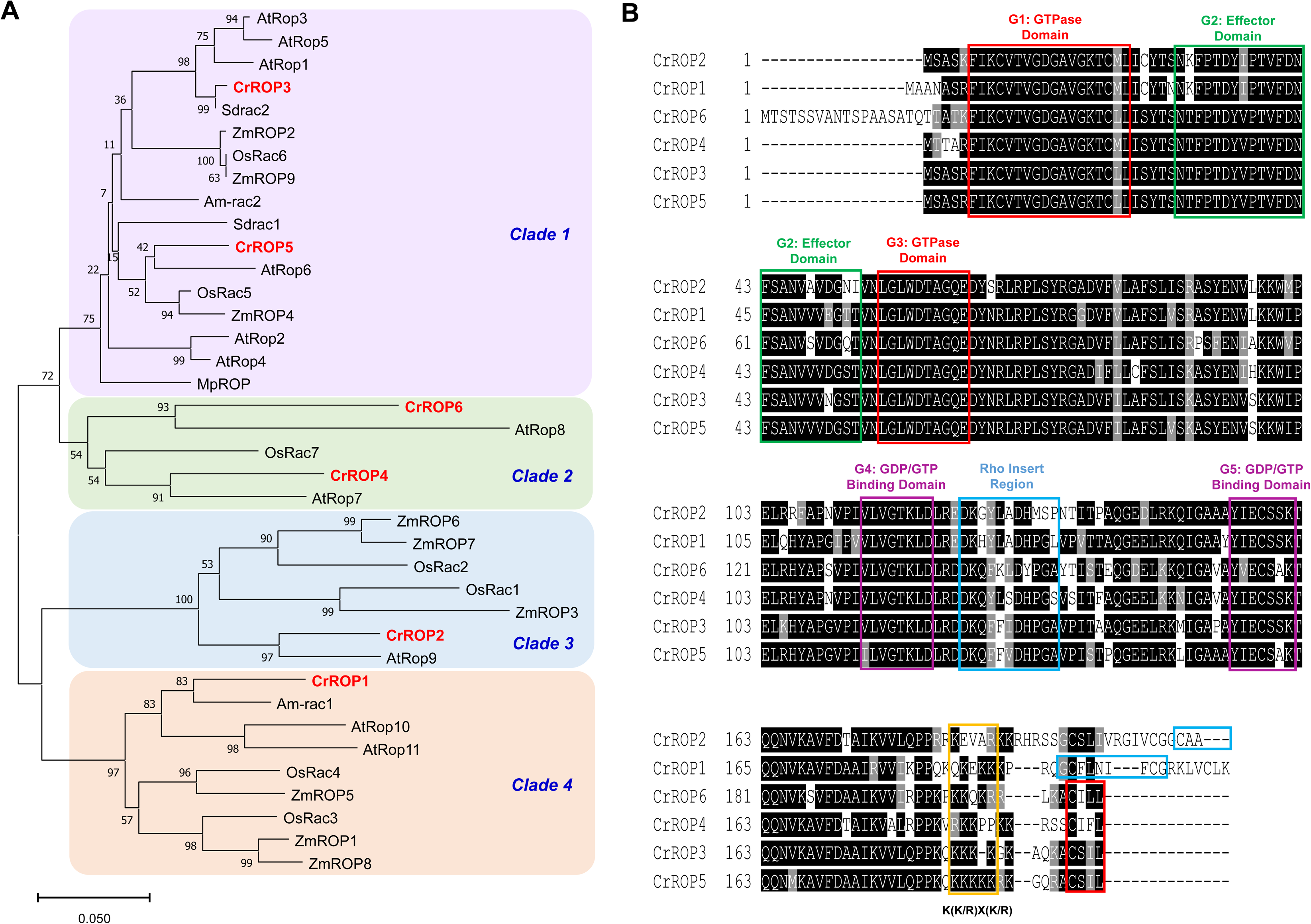
Sequence analyses of CrROPs. (A) Phylogenetic relationship of CrROPs with ROPs of other plants. The tree was constructed by neighbor-joining method with bootstrap value of 1000 runs using MEGA11 software. Accession numbers of sequences used to construct the tree are provided in Table S1. (B) Multiple sequence alignment of CrROPs was performed using Clustal W (Thompson et al., 1994) with default parameters through EMBnet (http://www.ch.embnet.org/software/ClustalW.html). Black and gray shadings was done with BOXSHADE 3.21 (http://www.ch.embnet.org/software/BOX_form.html) in which amino acids in black indicate conserved residues in all the six sequences, whereas resides in gray are similar in at least two of the sequences shown. Dashes indicate gaps inserted for optimal alignment. The conserved regions are marked with different coloured boxes and the nuclear localization signal (NLS) (K-K/R-x-K/R) is underlined.

Multiple sequence alignment of CrROP amino acid sequences revealed that they all possess five highly conserved G-Box motifs (G1-G5) that are characteristic of proteins belonging to Ras superfamily (Figure 2B). The GTPase domain contains G1 (GAVGKT) and G3 (DTAG) motifs, and the GTP/GDP binding domain contains G4 (VGTKL) and G5 (ECSS) motifs, whereas G2 (FPTDYVP) motif is part of effector domain (Figure 2B). In addition to conserved G-box motifs, ROPs contain hyper-variable C-terminus region, which determines the type of PTM. CrROP3, CrROP4, CrROP5, and CrROP6 consisted of conserved CxxL (C, cysteine; L, leucine; x, aliphatic amino acid) motif at the C-terminus and belonged to type-I ROPs, which undergo either farnesylation or geranylgeranylation. Among the four CrROPs, there was some variation in the types of aliphatic amino acids present in the “CxxL” motif. While CrROP3 and CrROP5 possessed the “CSIL” motif, CrROP4 and CrROP6 contained the “CIFL” and “CILL” motifs, respectively. Whereas CrROP1 and CrROP2 possessed FCGRKLVCLK and CAA motifs at the C-terminus, respectively, and belonged to type-II ROPs that undergo *S*-acylation (Figure 2B). Besides, the C-terminus of all six CrROPs contained a polybasic canonical nuclear localization signal sequence (K-K/R-X-K/R), in which the basic residue K is predominant with CrROP5 having the highest and CrROP1 with the least number of KR residues (Figure 2B).

### Spatio-temporal expression analysis of *CrROPs* in *C. roseus*

To determine the spatio-temporal gene expression of *CrROPs*, their transcript levels were assessed across different tissues such as leaf, flower, bud, silique, shoot, and root, by reverse transcriptase-quantitative polymerase chain reaction (RT-qPCR). The expression level of each *CrROP* in different tissues was calibrated with reference to its expression level in root, which was set to 1. All six *CrROP*s were expressed in different tissues, albeit at varying levels. However, *CrROP3* and *CrROP5* exhibited highest overall expression in aerial tissues compared to the other four *CrROP* candidates. Moreover, both *CrROP3* and *CrROP5* had highest expression in leaves, a very important site for MIA production in *C. roseus*. *CrROP3* exhibited the highest expression in leaves (21 fold), followed by buds and flowers (∼ 6 fold in both), and it showed a basal level of expression in roots, shoots, and siliques (Figure 3A). On the other hand, *CrROP5* had a highest but similar expression level in both leaves and flowers (∼ 25 fold), followed by buds (14 fold) and siliques (11 fold), with the least expression in roots and shoots. Among the others, *CrROP1* showed basal and similar expression levels in most of the tissues, while *CrROP2* exhibited the highest expression in buds (∼ 5 fold), followed by siliques, leaves, and roots, with the least expression in flowers and shoots (Figure 3A). Whereas *CrROP4* showed the highest expression in shoots (5 fold), with more or less similar transcript levels (ranging from ∼ 1 to 2 fold) in all other tissues, *CrROP6* had the highest expression in flowers (∼ 5 fold) and shoots (∼ 2.5 fold), with least and similar expression levels in other tissues (Figure 3A). In addition, a comparative expression analysis was conducted between different *CrROPs* by calibrating against *CrROP4*, which was set to 1 as it had the lowest expression level. The analysis revealed that among the six *CrROPs*, *CrROP3* and *CrROP5* exhibited relatively higher expression in most tissues with CrROP5 exhibiting overall higher expression than CrROP3 (Figure S2). This was evident even in *in silico* analysis of gene expression derived from MPGR dataset (Figure S3A). Further, to determine whether *CrROPs* are responsive to phytohormone methyl jasmonate (MeJA), a key elicitor of several specialized metabolic pathways, RT-qPCR analysis was performed. Geraniol synthase (CrGES), a key MIA pathway gene which is known to be induced by MeJA was used as a positive control. The results showed that unlike *CrGES* transcripts which were significantly induced, all *CrROPs* exhibited either no change or slight changes in their expression levels in response to MeJA (Figure S4). Even, *in silico* analysis using FPKM values derived from MPGR database showed no induction of *CrROP* expression in seedlings at different days interval (Figure S3B).

**Figure 3.**
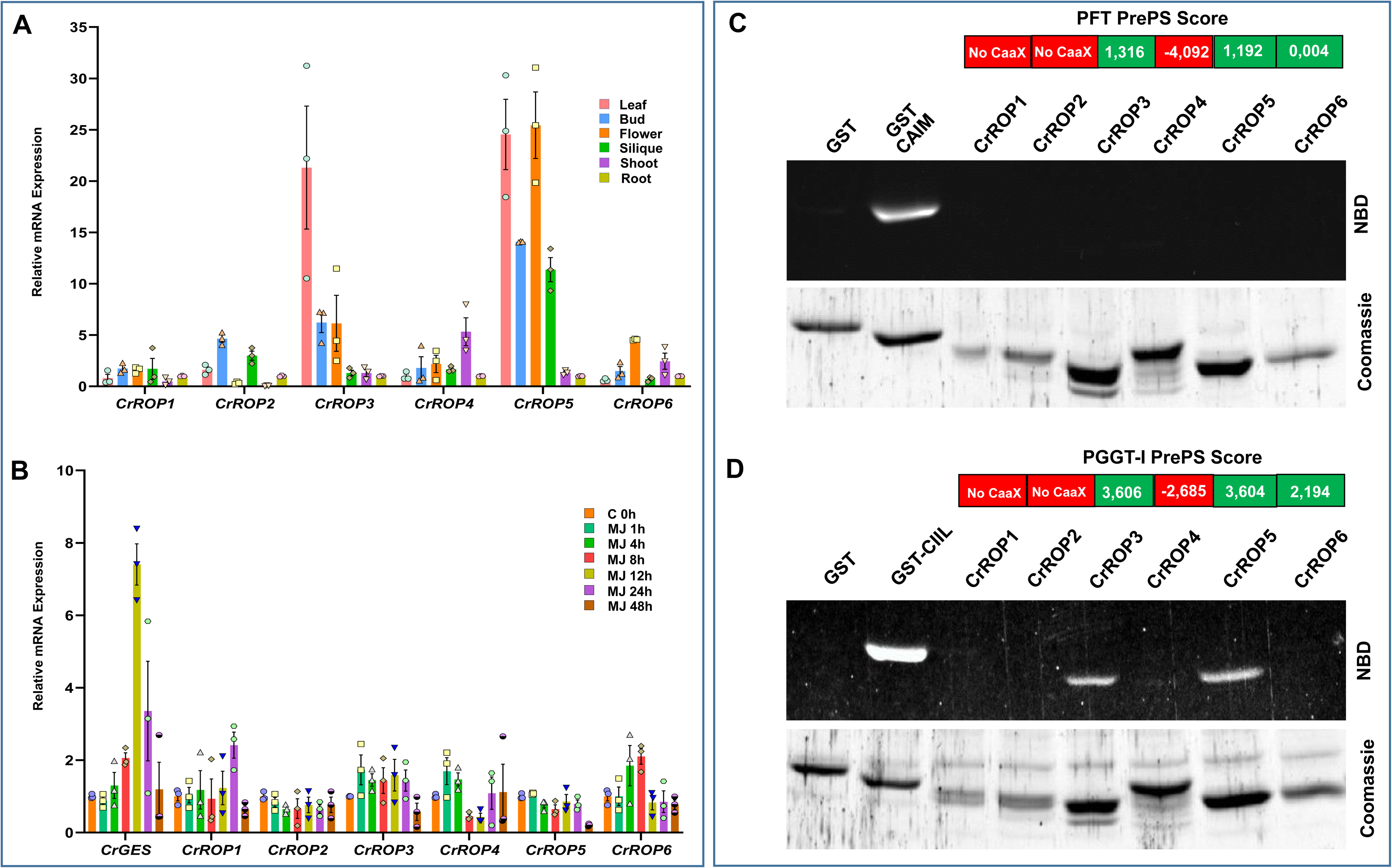
Spatio-temporal gene expression analysis and protein prenylation assay. (A) Relative transcript abundance of *CrROPs* in different tissues of *C. roseus*. Real time qPCR analysis was performed using total RNA extracted from leaf, flower, bud, silique, shoot and root. Expression levels of genes were normalized to the endogenous reference gene *CrN227* and are represented relative to root, which was set to 1. (B) Expression analysis of *CrROPs,* and *CrGES* in response to MeJA. Real time PCR analysis was performed using total RNA extracted from leaves treated with MeJA. Samples were collected at different time intervals. Statistics was performed with “GraphPad Prism 9” using two-way ANOVA: **, *P*=0.002; ***, *P*<0.001. Each data point represents the mean ± standard error (SE) of three independent experiments with three technical replicates. (C) *In vitro* farnesylation assay of CrROPs using recombinant *C. roseus* PFT. (D) In vitro geranylgeranylation assay of CrROPs using recombinant *C. roseus* protein geranylgeranyltransferase (PGGT-I). In vitro prenylation positive controls are GST fused with CAIM or CIIL motives for PFT or PGGT-I, respectively. NBD, NitroBenzoxadiazol fluorescent FPP (C) or GGPP (D) probe.

### CrROP3 and CrROP5 are geranylgeranylated by PGGT-I

The function of ROP GTPases depends on C-terminal stable prenylation or *S*-acylation. The presence of the canonical CaaL-box geranylgeranylation motif at the C-terminal end of CrROP3, CrROP4, CrROP5, and CrROP6 suggested that these proteins could be substrates for PGGT-I or PFT (Figure 2B). Moreover, among the six CrROPs, CrROP3, CrROP5, and CrROP6 were predicted to be geranylgeranylated using the PrePS prenylation prediction suite (http://mendel.imp.univie.ac.at/sat/PrePS). To experimentally determine whether CrROPs undergo prenylation, an *in vitro* prenylation assay was performed. Three different GST-tagged fusion proteins bearing a PFT motif (CAIM), a PGGT-I motif (CIIL), or an unprenylable motif (CWRL) were purified and used for the assay as controls. CrROPs were overexpressed in *E. coli* and the recombinant His-tagged proteins were purified by affinity chromatography (Figure S4A). The purified recombinant CrROP proteins were used for *in vitro* prenyltransferase assay. Incubation of purified CrROPs with fluorescent NBD-modified FPP and PFT, revealed that none of the six CrROPs were farnesylated in contrast to the positive control for farnesylation (i.e., GST-CAIM). On the other hand, incubation of fluorescent NBD-modified GGPP and PGGT-I showed that among the six CrROPs, only CrROP3 and CrROP5, having CSIL motif at the C-terminus, were geranylgeranylated similar to the positive control for geranylgeranylation (i.e. GST-CIIL). Further, activity assay using recombinant CrROPs in presence of GTP showed that all six CrROPs possessed functional GTPase activity (Figure S4B).

### CrROP3 and CrROP5 interact with CrPGGT-I

Given that CrROP3 and CrROP5 were geranylgeranylated, we hypothesized that CrPGGT-I could interact with these ROPs. *In silico* protein-protein interaction analysis using STRING software also supported this hypothesis wherein both type-I CrROP3 and CrROP5 were predicted to interact with CrPGGT-I, whereas no interaction was predicted between type-II CrROP2 and CrPGGT-I (Figure S5). To experimentally test our hypothesis and validate the STRING in-silico predictions, we assessed the interaction of CrROPs with CrPGGTI using the yeast two-hybrid system (Y2H). For conducting Y2H assays, we fused the activator domain (AD) of GAL4 at the N-terminus of CrROP2/3/5 (prey), while the GAL4 DNA binding domain (DBD) was fused at the N-terminus of CrPGGT-I (bait) (Figure 4A). Additionally, we prepared AD-CrROP3 and BD-CrROP5 constructs to investigate any potential interaction between the two CrROPs. Interaction assays revealed that yeast cells co-transformed with BD-CrPGGT-I (bait) and AD-EV (prey), AD-CrROP2 and BD-EV, AD- CrROP3 and BD-EV, and AD-CrROP5 and BD-EV showed growth in SD/-Trp/-Leu medium but did not grow on the SD/CLeu/CTrp/CHis/CAde (CLWHA) medium (Figure 4B&C).

**Figure 4.**
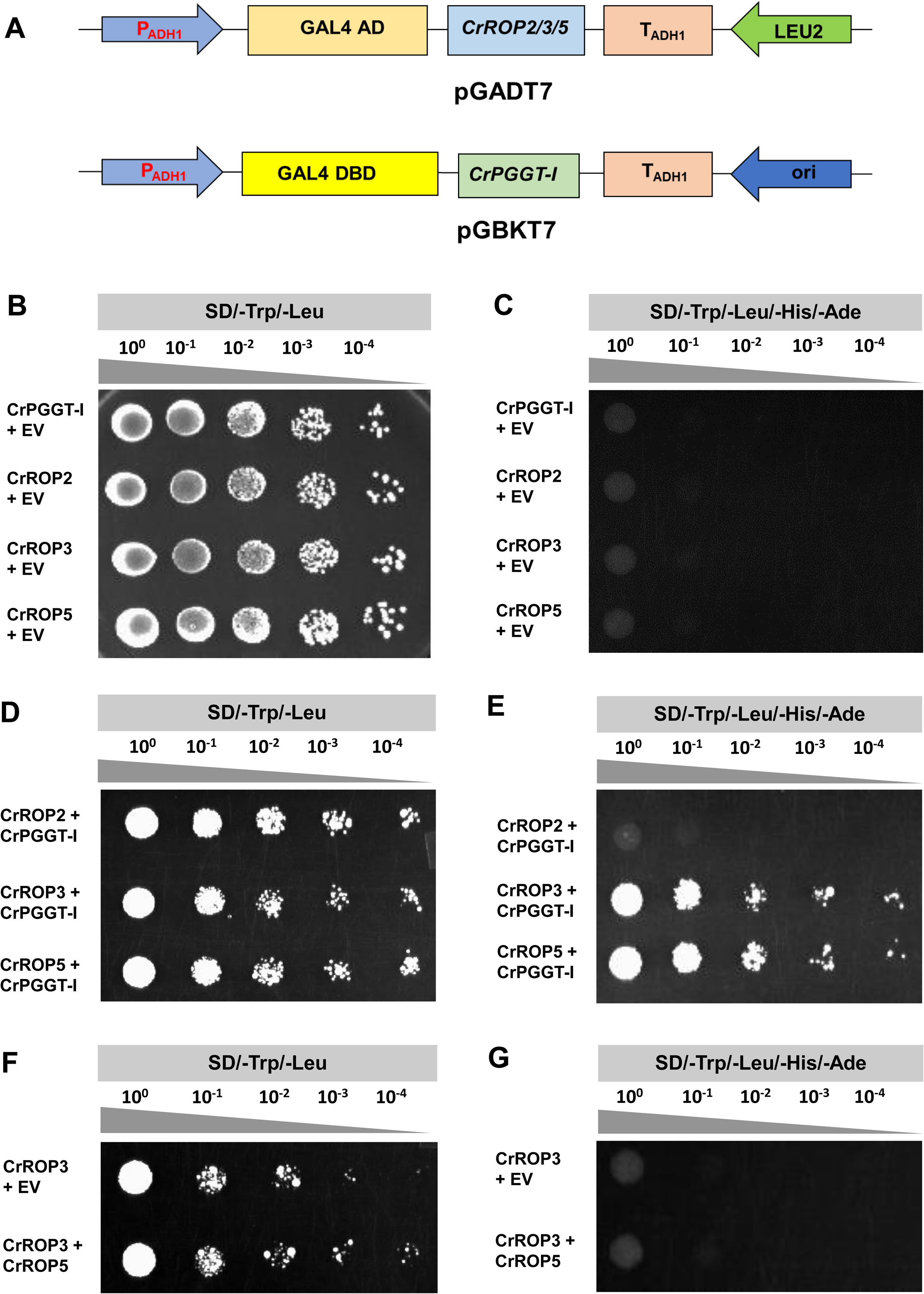
Yeast two-hybrid protein-protein interaction assay. (A&B) Vector maps of bait and prey constructs. The open-reading-frame (ORF) of CrROP2/3/5 were cloned in-frame with activation domain (AD) of pGADT7, and the ORF of CrPGGT-I was cloned in-frame with the DNA binding domain (DBD) of pGBKT7 vector. (C-H) Mating-based Yeast Two-Hybrid (Y2H) screening. CrROP2/3/5 were separately transformed into AH109 (MATa) yeast strain, which was then co-cultivated with the compatible mating type Y187 (MATα) yeast strain having CrPGGT-I or empty vector pGBKT7 (EV) to identify bait-interaction in the resulting diploid yeast cells. The transformants were plated using different dilutions on synthetically defined (SD) medium lacking leucin (Leu) and tryptophan (Trp) (SD/-Leu/-Trp) (C,E,G), and on SD/-Leu/-Trp medium lacking histidine and adenine (SD/-Leu/-Trp/-His/-Ade) (D,F,H). The plates were incubated for 3–5 days at 30C°C and observed for growth. Growth of yeast culture on SD/- Leu/-Trp/-His/-Ade indicates interaction of CrROP3 and CrROP5 with CrPGGT-I (F). Three independent colonies were tested per combination and a representative colony is shown.

However, yeast cells co-transformed with constructs of AD-CrROP3 and BD-CrPGGTI, and AD-CrROP5 and BD-CrPGGTI grew well on both SD-LW and SD-LWHA medium, whereas cells co-transformed with AD-CrROP2 and BD-CrPGGTI were unable to grow on SD-LWHA medium (Figure 4D&E). These results showed that both CrROP3 and CrROP5, which are type-I ROPs with the N-terminus “CSIL” motif, could interact with CrPGGT-I *in vivo*. However, CrROP2, which is a type-II ROP with the “GC…CG” motif, did not show interaction with CrPGGT-I (Figure 4D&E). No protein-protein interaction was seen between CrROP3 and CrROP5 in our Y2H analysis (Figure 4F&G).

### CrROP3 and CrROP5 are localized to nucleus, and CrROP2 is targeted to plasma membrane

ROP GTPases is generally considered to be localized to plasma membrane,, but they have also been reported to be localized to the nucleus, and cytosol (Chen et al., 2010; Ge et al., 2020; Han et al., 2022). Sequence analysis of CrROP2/3/5 revealed that the polybasic region of CrROP3 and CrROP5 harbors a K-K/R-x-K/R sequence, which is indicative of potential nuclear localization (Figure 2B). Whereas CrROP2 possessed least number of KR residues with CAA motif at the end of C-terminus (Figure 2B). As *in silico* prediction programs did not yield a consistent localization pattern for CrROP2/3/5 (Table S2), we assessed their subcellular localization experimentally by introducing green fluorescent protein (GFP) fusion protein (CrROP-GFP) into *C. roseus* cells. To ascertain the precise localization, each of the CrROP-GFP constructs was co-transformed with a plasmid expressing a specific marker for either plasma membrane (PM-mCH) or nuclear (nuc-mCH) localization. Analysis of fluorescence in co-transformed *C. roseus* cells by confocal microscopy revealed that CrROP2-GFP was predominantly localized at the cell’s periphery, indicating its localization in the plasma membrane (Figure 5A). Further, the overlapping signals emitted by the PM-mCH marker and the GFP signal from CrROP2-GFP provided clear evidence that CrROP2 is localized to the plasma membrane (Figure 5B&5C). In the case of CrROP3-GFP and CrROP5-GFP, the GFP signal was found to be exclusively associated with the nucleus (Figure 5E&I). The nuclear localization of CrROP3-GFP and CrROP5-GFP was further confirmed by the clear superimposition of the GFP signal with the signal emitted by the nuc- mCH marker (Figure 5F&5. Moreover, the localization of CrROP5-GFP was diffused in part of the nucleus, with a more pronounced localization in the nucleolus (Figure 5E,F&G).

**Figure 5.**
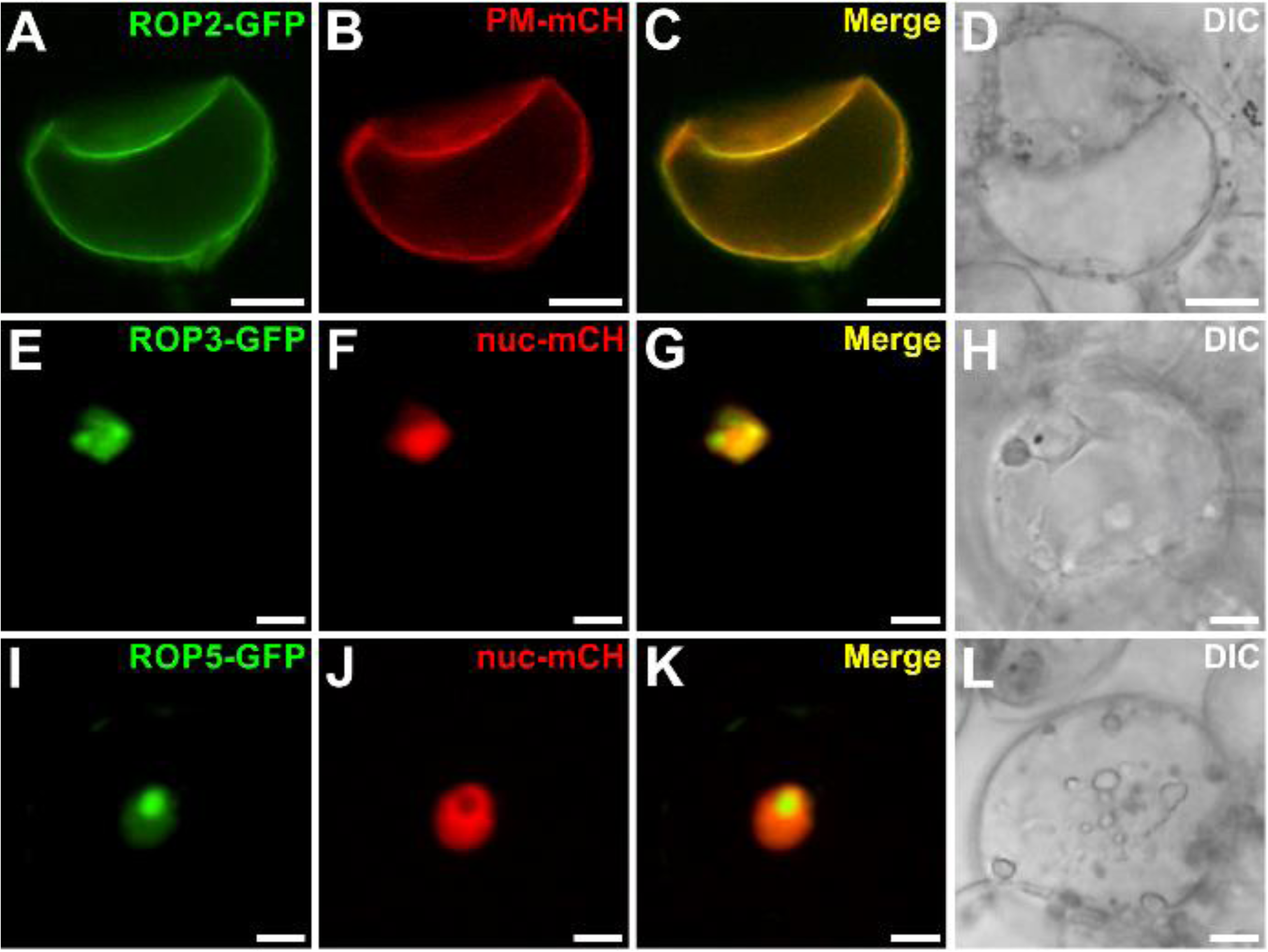
Analysis of subcellular localization of CrROP3, CrROP5 and CrROP2. Confocal laser scanning microscopy of *C. roseus* cells co-transformed with CrROP-GFP and PM-mCH (plasma membrane specific marker) or nuc-mCH (nuclear specific marker) expression constructs. In each subfigure, fluorescence detected from GFP and mCH fusion proteins is shown on the first and second vertical panels, respectively, whereas the third vertical panel shows merged GFP and mCH fluorescence. The fourth and the rightmost panel shows transmission image of cells.

### VIGS of *CrROP3* and *CrROP5* show their involvement in MIA biosynthesis

To discern the *in planta* role of *CrROP3* and *CrROP5* in MIA biosynthesis, the expression of these genes was down-regulated in *C. roseus* leaves by virus-induced gene silencing (VIGS) following the method reported previously (Kumar et al., 2015; Kumar et al., 2020). To prevent potential cross-silencing of *CrROP* isoforms, regions of *CrROP2*, *CrROP3* and *CrROP5* that showed the highest dissimilarity with other isoforms were utilized to generate VIGS constructs (Figure S6). In parallel, the pTRV2:*CrPDS* construct was used to silence the phytoene desaturase (PDS) gene, leading to a photobleached phenotype, which served as a visible marker for determining the timing to collect silenced leaf tissues (Figure 6A). The RT-qPCR expression analysis revealed that the silencing of *CrROP2/3/5* resulted in significant downregulation of their transcript levels in the range of 84-93% in newly emerging leaves of silenced plants compared to the empty vector (EV) control (Figure 6B). To evaluate the effect of these gene silencing on MIA biosynthesis, we measured the expression of genes related to MIA biosynthesis, such as TFs (*ORCA3, BIS2,* and *WRKY*), a kinase (*MPK3*), and pathway genes (*DXS, GES, G10H, AS, STR*, and *T16H*), which were previously shown to be affected in the *CrPGGT-I* silencing background (S. R. Kumar et al., 2020). RT-qPCR analysis revealed that the transcript levels of all analysed TFs, MPK3 and pathway genes were drastically declined by 40 to 92% in *CrROP3*-, and *CrROP5*-vigs leaf tissues compared with EV (Figure 6D&E). In contrast, samples derived from *CrROP2*-vigs plants showed no significant change in the expression of any of the above analysed genes related to MIA biosynthesis as compared to EV control (Figure 6C).

**Figure 6.**
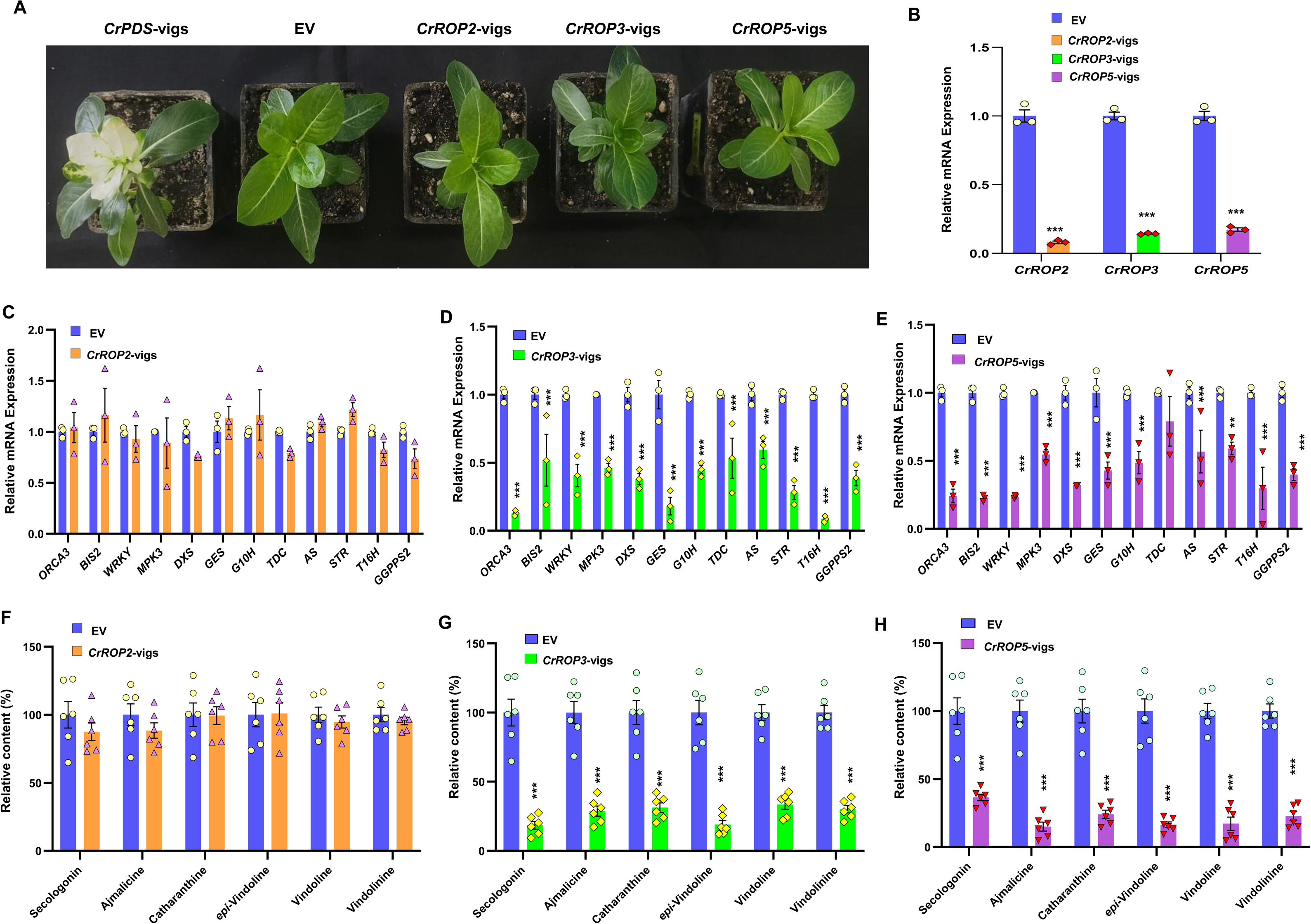
Effect of virus-induced gene silencing of *CrROP2/3/5* on MIA biosynthesis. (A). Representative images of plants infected with *Agrobacterium tumefaciens* harboring pTRV vectors and VIGS constructs. Images were taken 3 weeks after infection. (B) Reverse transcriptase-quantitative polymerase chain reaction (RT-qPCR) analysis showing the relative expression of CrROP2/3/5 in comparison to EV control. Effect of CrROP2/3/5 silencing on expression of TFs and pathway genes related to MIA biosynthesis (C,D,E) and major MIAs (F,G,H). Expression level of genes was normalized to *CrN227* endogenous control and was set to 1 in EV to determine the relative reduction in *CrROP*-vigs leaves. Levels of different metabolites are represented relative to EV control. The results shown are from three (qPCR) and six (metabolites) independent biological replicates with 3 and 2 technical replicates, respectively. Statistics was performed with “GraphPad Prism 9” using one-way ANOVA: ***, *P*<0.001. Error bars indicate mean ± SE.

Next, analysis of metabolites showed that specific silencing of *CrROP3* and *CrROP5*, which led to downregulation of genes related to MIA biosynthesis, resulted in significant reduction in the levels of secologanin (a key iridoid intermediate of MIA pathway) and downstream MIA (Figure 6G&H). The leaves of *CrROP3*-vigs plants contained 81.5% lower levels of secologanin than that EV leaves (Figure 6G). This was followed by a significant decrease in the levels of *epi*-vindolinine (81%), vindolinine (70%), vindoline (67.3%), catharanthine (68.8%), and ajmalicine (72.2%). Similarly, *CrROP5*-vigs leaves showed a drastic decline in secologanin (63.5%), followed by vindoline (82.8%), vindolinine (77.4%) and ajmalicine (85%) content as compared to EV control leaves (Figure 6H). Unlike *CrROP3* and *CrROP5* silencing, which resulted in a drastic reduction in MIA biosynthesis, *C. roseus* plants with suppressed *CrROP2* did not exhibit any significant decrease in the levels of either the iridoid intermediate secologanin or downstream MIA (Figure 6F).

### Overexpression of CrROP3 and CrROP5 enhances MIA biosynthesis

Suppression of *CrROP3* and *CrROP5,* and not *CrROP2,* negatively affected the MIA biosynthesis (Figure 6C-H). To further elucidate the effect of *CrROP2*, *CrROP3*, and *CrROP5* overexpression on MIA biosynthesis, the coding sequence of these genes was placed under the control of the cauliflower mosaic virus (CaMV) 35S promoter, and transiently overexpressed them individually in *C. roseus* by agroinfiltration. Analysis of *CrROP2*, *CrROP3*, and *CrROP5* expression showed an 18.6 fold, 12.82 fold, and 29.83 fold increase in their transcript levels, respectively. It was observed that despite an 18.6 fold increase in the transcript level of *CrROP2*, there was no significant difference in the expression level of genes related to MIA biosynthesis or in the levels of secologanin and MIA between 35S:*CrROP2* and EV control (Figure 7A&D). On the other hand, leaves overexpressing *CrROP3* and *CrROP5* exhibited a corresponding and significant enhancement in the expression of all TFs, MPK3, and most MIA pathway genes. The increase in gene expression ranged between 2.5-17.09 folds (except for *HMGR1* and *AS*, which did not exhibit a significant enhancement) with a maximum observed for *G10H* and a minimum but significant enhancement for *TDC* in *CrROP3* overexpressing samples (Figure 7B). Similarly, *CrROP5* overexpressing samples displayed a corresponding increased expression of genes which ranged between 6.60-35.46 folds (except for *HMGR1*, *TDC*, *AS*, and *T16H* which did not exhibit a significant enhancement) with a maximum observed for *STR* and a minimum *DXS2* (Figure 7C). Subsequent quantification of metabolites revealed a significant elevation in the levels of analyzed metabolites in both *CrROP3* and *CrROP5* overexpressing leaves compared with the EV control. The degree of enhancement in secologanin and MIA levels was consistent with the degree of overall increase in expression of genes related to MIA biosynthesis in *CrROP3* and *CrROP5* overexpressing samples. While *CrROP3* overexpressing samples showed 1.82 fold and ∼1.5 to 1.9 fold increased accumulation of secologanin and different MIA, respectively, *CrROP5* overexpression exhibited 2 to 3.7 fold increased accumulation of those metabolites (Figure 7E&F).

**Figure 7.**
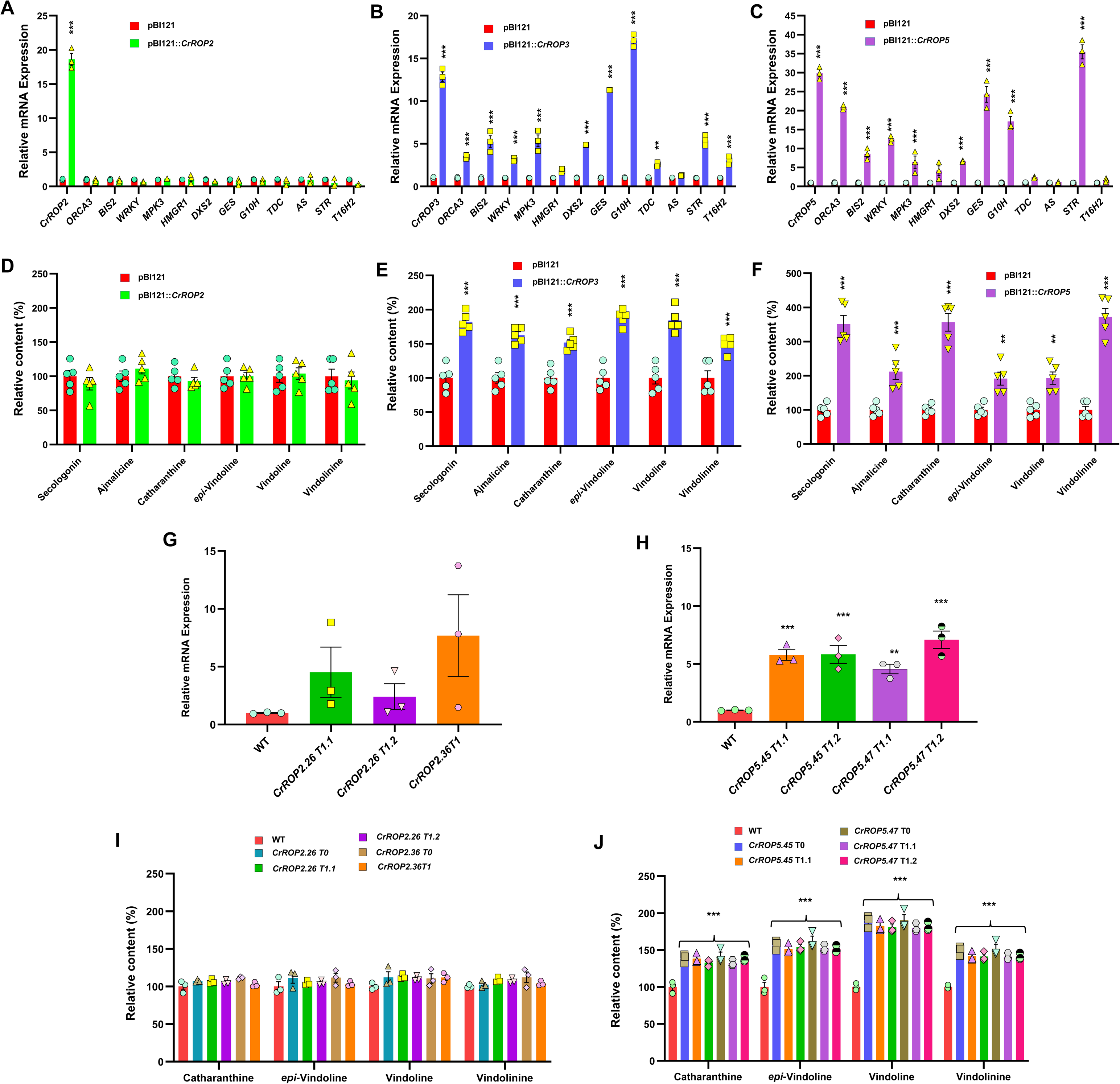
Overexpression of *CrROP2/3/5* and its effect on *C. roseus* MIA biosynthesis. qRT-PCR analysis of gene expression of *CrROPs,* TFs and pathway genes related to MIA biosynthesis in *C. roseus* agroinfiltrated with control (pBI121) and CrROP2/3/5 (A-C), and the corresponding effect on MIA accumulation (D-F). (G-H) Relative expression of CrROP2/5 in wild type (WT) and transgenic T1 lines of *C. roseus*. (I-J) Relative levels of major MIAs in WT and transgenic lines. Expression levels of genes were normalized to the endogenous reference gene *CrN227* and are represented relative to the pBI121 control (EV) control, which was set to 1. Relative amounts of MIAs were analyzed by HPLC and are expressed in % relative to pBI121 (D-F) and WT (I-J) controls. The results shown are from three to five biological replicates. Statistics was performed with “GraphPad Prism 9” using two-way ANOVA: **, *P*=0.002; ***, *P*<0.001. Error bars indicate mean ± SE.

To further substantiate the results obtained in transient overexpression studies, transformed *C. roseus* plants overexpressing *CrROPs* were generated through tissue culture-independent Agrobacterium-mediated *in planta* transformation as described earlier (S. R. Kumar et al., 2020).. For both *CrROP2* and *CrROP5* two independent transformed plants (T_0_) were obtained, however, despite repeated efforts, *CrROP3* overexpressing plants could not be generated. To obtain T_1_ plants, seedlings were obtained by germinating seeds of T_0_ plants, and PCR-screened to confirm the transgenic nature. For *CrROP2* two T_1_ plants for line *CrROP2.26* and one plant for line *CrROP2.36* were obtained whereas two T1 plants for each line *CrROP5.45* and *CrROP5.47* were confirmed by pCR screening (Figure S7). The transgenic lines showed increased expression of the corresponding introduced gene which ranged from 2.4 fold to 7.6 fold for *CrROP2* and 4.5 fold to 7 fold for *CrROP5* compared to the WT control plants (Figure 7G&H). Subsequent metabolite analysis revealed that stable overexpression of *CrROP5* resulted in an enhanced accumulation of MIA (1.4 to 1.9 fold), whereas overexpression of *CrROP2* did not show any significant change in the MIA levels compared to the WT (Figure 7I&J). These findings were consistent with the results obtained from the transient overexpression studies involving *CrROP2* and *CrROP5* (Figure 7D&F).

### Removal of CSIL-motif in CrROP3 and CrROP5 fails to enhance MIA biosynthesis

CSIL-motif containing CrROP3 and CrROP5 proteins were not only geranygeranylated by CrPGGT-I but also interacted with it (Figure 3D & 4E). Moreover, both CrROP3 and CrROP5 showed *in planta* functional role in modulation of MIA biosynthesis (Figure 6&7). This prompted us to investigate whether the CSIL motif is essential for CrROP3 and CrROP5 to exert a modulatory effect on MIA biosynthesis. Therefore, we generated Δ*CrROP3* and Δ*CrROP5* mutants lacking 12 nucleotides that encode amino acids corresponding to the C-terminal “CSIL” motif (Figure 8A&B). To validate the *in planta* function, constructs for wild-type, and mutant *CrROP3* and *CrROP5* along with vector control were individually transformed into *C. roseus* leaves by Agroinfiltration. Analysis of transcript levels showed that both wild-type, and mutant *CrROP3* and *CrROP5* had elevated expression ranging from 6.56 fold to 8.6 fold compared to the EV control (Figure 8C&D). While this increase in transcript levels significantly enhanced the accumulation of MIA by 1.5 fold to 2.17 fold in the wild-type background compared to the EV control, there was no effect on the accumulation of any of the analyzed MIA in leaves overexpressing Δ*CrROP3* or Δ*CrROP5* (Figure 8E&F). Moreover, the level of analyzed MIA in leaves infiltrated with Δ*CrROP3* or Δ*CrROP5* was almost identical to the level found in leaves transformed with the EV control (Figure 8E&F).

**Figure 8.**
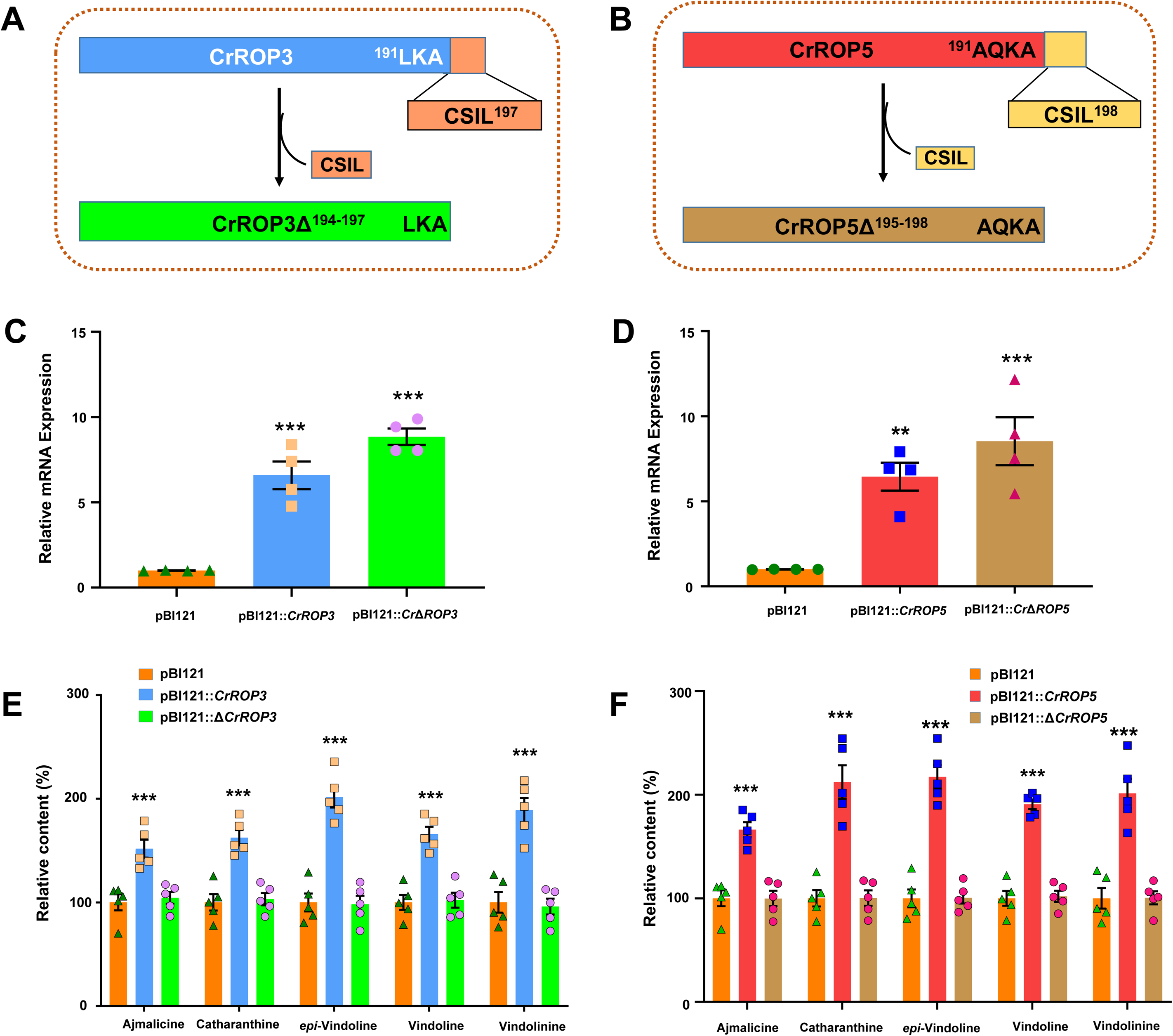
Effect of *CrROP3/5* mutants lacking CSIL motif on MIA biosynthesis. (A&B) Schematic representation of *CrROP3* and *CrROP5* showing CSIL motif. Relative expression of Δ*CrROP3* and Δ*CrROP5* in *C. roseus* leaves agroinfiltrated with pBI121, pBI121::*CrROP3/5,* and pBI121::Δ*CrROP3/5* constructs (C&D) and the corresponding effect on accumulation of major MIAs (E&F). Expression levels of genes were normalized to the endogenous reference gene *CrN227* and are represented relative to the pBI121 control (EV) control, which was set to 1. Levels of MIAs were analyzed by HPLC and are expressed in % relative to pBI121 control. Statistics was performed with “GraphPad Prism 9” using two-way ANOVA: ***, *P*<0.001. Error bars indicate mean ± SE.

## DISCUSSION

It has been demonstrated that disruption of geranylgeranylation has a negative impact on MIA biosynthesis in *C. roseus* and sesquiterpene capsidiol production in tobacco (Courdavault et al., 2009; Huchelmann et al., 2014; Kumar et al., 2020). Moreover, the availability of the geranylgeranyl precursor GGPP, necessary for geranylgeranylation, has been shown to play critical role in regulation of *C. roseus* MIA biosynthesis (Courdavault et al., 2005; Courdavault et al., 2009; Kumar et al., 2018; Kumar et al., 2020). These studies highlight the significance of geranylgeranylation and indicate the involvement of geranylgeranylated proteins in the regulation of specialized metabolism in plants. Given this information, and considering that ROP proteins are known to undergo prenylation (Sorek et al., 2010; Sorek et al., 2011), and that overexpression of a ROP from *S. dulcis* (*Sdrac2*) enhanced atropine content in *A. belladonna* (Asano et al., 2013), we hypothesized specific geranylgeranylated ROPs may play a role in the regulation of MIA biosynthesis in *C. roseus*. Hence, the current study aimed to conduct a detailed investigation on the role of ROPs in *C. roseus* MIA biosynthesis.

Plants contain a varying number of ROPs, ranging from one in *Marchantia polymorpha* to four in *Physcomitrella patens* among lower plants, and from four in *Pinus taeda* (loblolly pine) to eleven in Arabidopsis and thirteen in poplar among higher plants (Rong et al., 2022; Fowler, 2010; Feiguelman et al., 2018). *C. roseus* has six ROP candidates which are distributed among 4 of the 8 chromosomes in the genome (Figure S1B). All six CrROPs possessed a conserved G-domain, a backbone of five G-boxes characteristic of small GTPases, which is involved in GTP-binding, hydrolysis, and interaction with target proteins (Feiguelman et al., 2018). Moreover, all six CrROPs exhibited *in vitro* GTPase activity, confirming that the chosen candidates are indeed small GTPases (Figure S4). Phylogenetically, CrROPs displayed significant similarity to ROPs of *Arabidopsis* and other land plants, and they could be categorized into four distinct clades (Figure 2A). CrROP1 belongs to clade-4 and shares a high sequence similarity with AmRac1, and also with AtROP10 and AtROP11, which have been shown to be involved in suppression of ABA signaling, secondary cell wall patterning, root hair growth, and inhibition of endocytosis (Choi et al., 2014; Xin et al., 2005; Feiguelman et al., 2018), CrROP2 belongs to clade-3 and exhibited high sequence similarity to AtROP9, which is known to regulate ABA and auxin signaling, and migration of pollen nuclei (Li et al., 2013; Nibau et al., 2013; Choi et al., 2014). Whereas CrROP4 and CrROP6 clustered in clade-2 together with AtROP7 and AtROP8, which have pivotal roles in vascular development and pollen nuclei movement (Wang et al., 2017; Kawashima et al., 2014). On the other hand, CrROP3 and CrROP5 were grouped under clade-1, which also includes Arabidopsis AtROP2, AtROP4, and AtROP6 that regulate root hair initiation, ROS production, phytochrome-mediated light responses, ABA signaling, and SA-mediated pathogen responses (Poraty-Gavra et al., 2013; Feiguelman et al., 2018; Jones et al., 2002; Molendijk, 2001). Moreover, CrROP3 and CrROP5 clearly clustered together with AmRac2 and SdRac2, which positively affect δ-guaiene synthase expression and atropine accumulation, respectively, suggesting that the ROPs from this clade could also be involved in the regulation of specialized metabolic pathways. Notably, clade-1 also includes the sole ROP of *M. polymorpha*, a bryophyte (Rong et al., 2022), suggesting that the ROP candidates in land plants first originated within this clade.

PTMs of ROPs represent an additional regulatory layer that facilitates the integration of functional modulation and subcellular localization, enabling precise spatiotemporal control of GTPase activity. Moreover, tight regulation of different types of PTMs is crucial to ensure accurate and appropriate GTPase signaling (Lemichez et al., 2020; Olson, 2018). ROPs are divided into two types based on the “motif” present at the C-terminal hyper variable region that dictates the type of PTM (Winge et al., 2000). Type-I ROPs possess “CaaL” motif and undergo prenylation, whereas type-II ROPs have either GC-CG or CAA motifs at the C-terminus and undergo *S-*acylation (Lavy & Yalovsky, 2006; Sorek et al., 2011; Feiguelman et al., 2018). In type-I ROPs, the Cys residue undergoes protein geranylgeranylation by the C_20_ isoprenyl lipid geranylgeranyl in a reaction catalyzed by PGGT-I, which can recognize the C-terminal CaaL motif (Figure 3C&D)(Sorek et al., 2011; Bao et al., 2022). The presence of the CaaL motif at the C-terminal end of CrROP3, CrROP4, CrROP5 and CrROP6 all of which belong to type-I ROPs suggested that these proteins are possible substrates of CrPGGT-I. However, further experimental evidence showed that only CrROP3 and CrROP5 were geranylgeranylated by CrPGGT-I, and none of the six CrROPs were farnesylated by PFT (Figure 3C&D). These results establish that CrROP3 and CrROP5 possessing functional “CSIL” motif efficiently get geranylgeranylated *in vitro*. Therefore, activation of CrROP3 and CrROP5 via prenylation by CrPGGT-I could be a critical process at the tissue level to control MIA biosynthesis in *C. roseus*. Interestingly, the transcripts of CrROP3 and CrROP5 that were geranylgeranylated exhibited higher expression in most tissues, albeit at different levels, indicating their role in different processes. Moreover, expression of *CrROPs* was not affected much in response to treatment with MeJA, a known inducer of MIA biosynthesis in *C. roseus* (Rischer et al., 2006; Oudin et al., 2007). The uninduced expression of *CrROPs* was similar to the expression of *CrGGPPS2* and *CrPGGT-I* in response to MeJA (Kumar et al., 2020; Courdavault et al., 2009). This indicated that transcripts of *CrROP3* and *CrROP5* co-expressed with the transcripts of their related upstream genes *CrPGGT-I* and *CrGGPPS2,* which have been shown to regulate MIA biosynthesis in *C. roseus* in previous studies (Courdavault et al., 2005; Courdavault et al., 2009; Kumar et al., 2020).

Though a coordinated role for CrGGPPS2 and CrPGGT-I in MIA biosynthesis has been proposed (Kumar et al., 2020), a direct connection between CrPGGT-I and CrROPs remains unclear. It is a fact that CrPGGT-I catalyzes the transfer of a C_20_ isoprenoid moiety to downstream proteins, thus facilitating their subcellular localization and protein-protein interactions (Sorek et al., 2011; Running, 2014; Bao et al., 2022). Interestingly, both CrROP3 and CrROP5, which possess the “CSIL” motif, were found to interact with CrPGGT-I, the enzyme responsible for protein geranylgeranylation. On the other hand, CrROP2, lacking the “CaaL” motif, was neither geranylgeranylated nor showed any interaction with CrPGGT-I (Figure 4). In a recent study, a type-I GhROP3 having the C-terminal “CAFL” motif was found to interact with GhGGB (PGGT-I) resulting in negative regulation of ABA signaling in cotton (Hu et al., 2022). In addition to PTM, subcellular localization is a fundamental aspect of a protein’s function, as it determines where and how a protein exerts its molecular actions (Pan et al., 2021). For ROPs, their localization to specific cellular compartments is essential for their proper functioning and interaction with downstream effectors. It is known that ROP proteins are synthesized as soluble proteins and undergo successive lipid modification involving prenylation or palmitoylation, which anchor the proteins to the membrane (Sorek et al., 2011; Lavy & Yalovsky, 2006; Feiguelman et al., 2018). Several studies have demonstrated that both type-I and type-II ROPs have plasma membrane localization where they trigger signaling, but for the type-I ROPs/RACs, localization could be found also in the cytoplasm and nucleus (Sorek et al., 2010; Chen et al., 2010; Sorek et al., 2011; Feiguelman et al., 2018; Han et al., 2022; Xiang et al., 2023). Interestingly, most ROPs contain a canonical nuclear localization signal (NLS) sequence [K(K/R)X(K/R)] (Chen et al., 2010). Both CrROP3 and CrROP5 were found to contain a NLS sequence, and were localized in the nucleus. In contrast, ROP2 did not possess an NLS and was instead localized to the plasma membrane (Figure 5). Sequence analysis revealed that both CrROP3 and CrROP5 belong to the type-I ROP category and were localized exclusively within the nucleus (Figure 5). Nevertheless, it is plausible that these proteins might also exhibit cytosolic or plasma membrane localizations intermittently, attributed to the protein’s functional cycles regulated by cytoplasmic or plasma membrane shuttling mechanisms, as previously shown for other ROP proteins (Yalovsky, 2015; Hodge & Ridley, 2016). These results suggest that the interaction of CrROP3 or CrROP5 with CrPGGT-I facilitates their geranylgeranylation and subcellular localization, potentially influencing downstream signaling and leading to the regulation of MIA biosynthesis.

To examine this, we adopted a reverseCgenetics VIGS approach to silence the *in planta* expression of *CrROP3* and *CrROP5*, as well as *CrROP2* that served as a control. Interestingly, specific silencing of *CrROP3* and *CrROP5*, both of which end with the CSIL C-terminal motif, resulted in drastic decrease in the transcript levels of TFs (*ORCA3, BIS, WRKY, MPK3*) and genes (*DXS, GES, G10H, STR, T16H, AS* and *TDC*) related to MIA biosynthesis (Figure 6D&E). This reduced transcript level was accompanied by a significant reduction in the levels of secologanin and MIA (Figure 6G&H). The effect on ajmalicine, catharanthine and vindoline accumulation was stronger in the *CrROP5*-silenced plant, consistent with the higher expression level of *CrROP5* as compared with *CrROP3* in *C. roseus* leaves (Figure 3A; Figure 6G&H). However, no effect was seen in the growth and morphology of *CrROP3*-vigs and *CrROP5*-vigs plants. In contrast, silencing of *CrROP2* did not exhibit any change in the expression level of TFs and genes related to MIA biosynthesis, and also in the level of metabolites (Figure 6C&F). This was in agreement with the previous reports that silencing of genes that are involved in geranylgeranylation (PGGT-I) or inhibition/suppression of geranylgeranylation negatively impacts MIA biosynthesis in *C. roseus* and capsidiol production in tobacco (Kumar et al., 2020; Courdavault et al., 2009; Huchelmann et al., 2014). Correspondingly, the reverse effect of silencing was observed upon transient overexpression of *CrROP3* and *CrROP5* (Figure 7). The expression of most of the TFs and genes that showed reduced expression in the silencing background was upregulated, resulting in corresponding enhancement in the accumulation of secologanin and MIA (Figure 7). Moreover, stable overexpression of CrROP5 also enhanced the MIA accumulation in *C. roseus*, further confirming the results obtained in transient overexpression. In fact, the effect of *CrROP3* and *CrROP5* overexpression was similar to the effect observed in silencing and overexpression of *CrGGPPS2*, which provides GGPP precursor for CrPGGT-I (Kumar et al., 2020). In agreement with our results, overexpression of a type-I ROP from *S. dulcis* Sdrac2 enhanced the biosynthesis of tropane alkaloid atropine in *A. bellodona* (Asano et al., 2013), indicating a possibility of a similar scenario in other alkaloid producing plants. Although the transcript level of *CrROP2* was significantly enhanced in both transient and stable overexpression backgrounds, it had no effect on either the transcript levels of TFs and genes related to MIA biosynthesis or the levels of secologanin and MIA (Figure 7). Based on these results, it can be construed that CrROP3 and CrROP5 act as *bona fide* signaling proteins and play a role in triggering the signaling cascade leading to the enhanced accumulation of MIA.

VIGS and overexpression experiments clearly established that among the three CrROPs (CrROP2, CrROP3, and CrROP5), only CrROP3 and CrROP5, which possessed the “CSIL” motif, affected MIA biosynthesis (Figure 6 & 7). Moreover, both CrROP3 and CrROP5 were geranylgeranylated by CrPGGT-I and interacted with CrPGGT-I (Figure 3D & Figure 4). These results clearly highlight the importance of the “CSIL” motif for geranylgeranylation, which in turn modulates MIA biosynthesis. Earlier investigations have also indicated that type-I ROPs, featuring a “CaaL” motif at the C-terminus, undergo geranylgeranylation (Sorek et al., 2011). Further, it was reported that the presence of Arg/Lys-rich polybasic domain in close proximity to the “CaaL” motif enhances the rate of geranylgeranylation (Caldelari et al., 2001). Indeed, CrROP3 and CrROP5 contain a lysine-rich region at the C-terminal hypervariable region, whereas this feature is not as evident in CrROP4 and CrROP6 (Figure 2B). This could also be a reason for CrROP3 and CrROP5 getting geranylgeranylated whereas CrROP4 and CrROP6 are not gernaylageranylated despite of having “CaaL” motif. To further substantiate the role of the “CSIL” motif in CrROPs on MIA biosynthesis, mutant ΔCrROP3 and ΔCrROP5 devoid of the CSIL motif were overexpressed. Unlike wild-type CrROP3 and CrROP5, which as shown before enhanced MIA biosynthesis, the mutants showed no effect on the level of MIA accumulation. This indicated that the “CSIL” is necessary for protein geranylgeranylation of CrROP3 and CrROP5, which in turn modulate MIA biosynthesis (Figure 8E & F). Therefore, it can be inferred that the overexpression of mutants lacking the “CSIL” motif resulted in non-prenylatable ΔCrROP3 and ΔCrROP5, which might not have been geranylgeranylated *in vivo* by CrPGGT-I at a sufficient level. Hence, the mutant ΔCrROP3 and ΔCrROP5 proteins could be possibly mis-localized and unable to initiate a signaling mechanism that transcriptionally upregulates MIA biosynthesis indicating the critical function of “CSIL” motif.

The fact that CrROP3 and CrROP5 interact with CrPGGT-I, get geranylgeranylated, localize to the nucleus, and positively affect MIA biosynthesis indicates that the modulation of gene expression and MIA accumulation occurs in the nucleus. Recently, several tomato ROPs have been shown to be localized in the nucleus; however, their exact roles have not been determined (Wang et al., 2022). Taking cues from these observations, as well as results from this study, we checked the interaction of CrROP3 and CrROP5 with TFs related to MIA biosynthesis that were affected in CrROP3 and CrROP5 silencing and overexpression backgrounds. None of the tested TFs (*WRKY1, BIS2, MPK3, MYC2, ORCA3*) showed any interaction with CrROP3 or CrROP5 (Figure S8). In another study, it was demonstrated that a tobacco ROP (NtRHO1) containing the “CSIL” motif is localized in the nucleus, where it interacts with a WRKY TF to negatively regulate defense against tobacco mosaic virus (Han et al., 2022). The absence of interaction between CrROP3 and CrROP5 with any of the tested TFs suggests the potential existence of an alternative mechanism wherein nuclear-localized ROPs might function together with nuclear-localized ROP-associated effectors, such as GTPase-activating proteins (GAPs) and guanine nucleotide exchange factors (GEFs). This scenario has been demonstrated in Human Embryonic Kidney (HEK) cells, where RhoA engages with RhoA-GAP (DLC1) and RhoA-GEFs (Net1 and Ect2) within the nucleus (Dubash et al., 2011).

In conclusion, we have uncovered a novel role of ROPs and demonstrated that specific CrROPs, namely CrROP3 and CrROP5, which contain the geranylgeranylation motif “CSIL,” play a positive regulatory role in *C. roseus* MIA biosynthesis. Thus, our data suggest that geranylgeranylation/ROP module is essential signaling component of MIA biosynthesis in *C. roseus* and that CrROP3 and CrROP5 function downstream of CrPGGT-I and upstream of JA signaling for modulation of MIA biosynthesis (Figure 9). These findings shed light on the significance of ROP GTPases in controlling the biosynthesis of specialized metabolites in plants. Further exploration of the downstream effectors of ROPs, which is a subject of our ongoing investigation, would provide deeper insights into the underlying mechanism of specialized metabolic pathway regulation.

**Figure 9.**
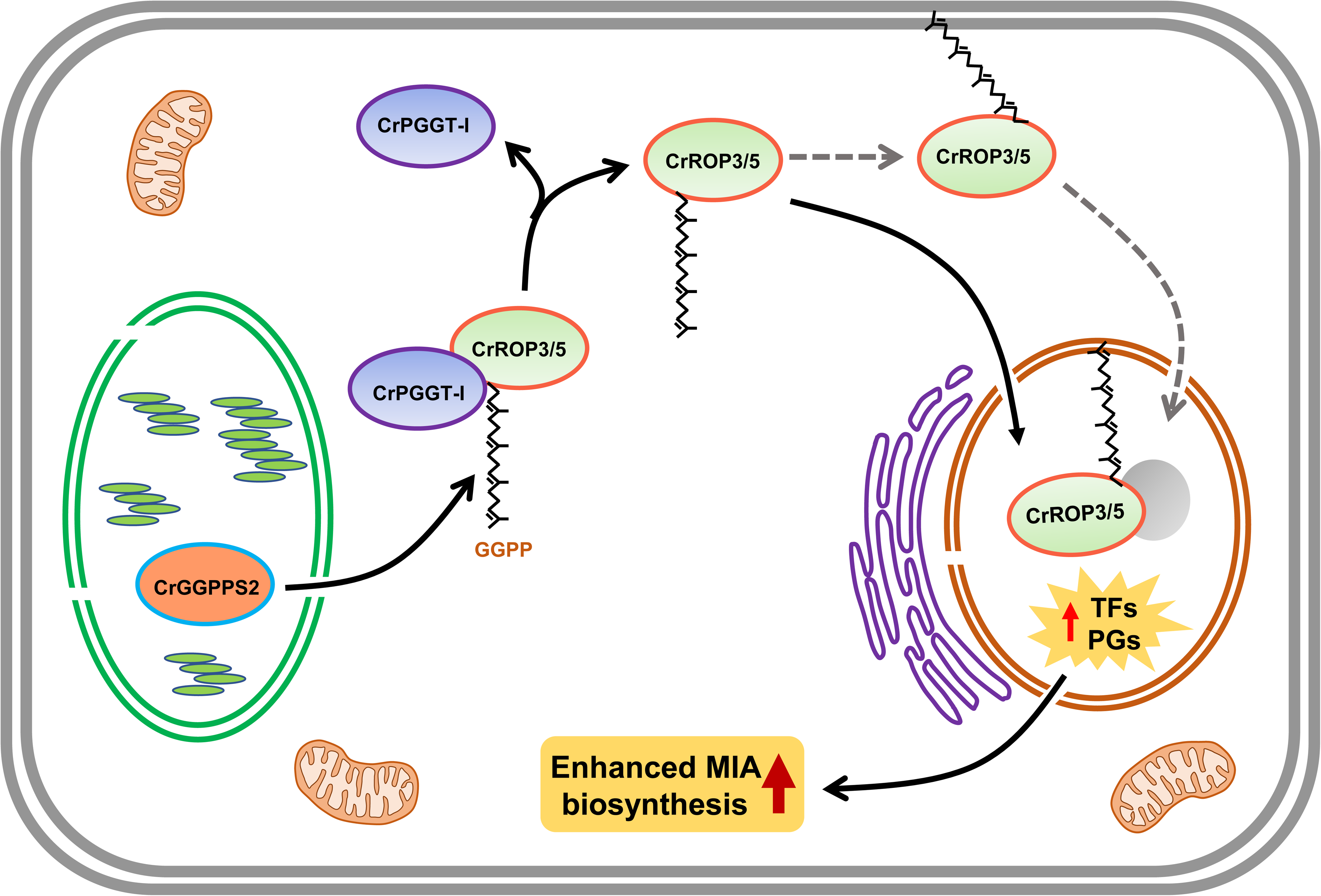
Proposed model showing the role of CrROP3/5 in modulation of terpene indole alkaloids accumulation in periwinkle.

## MATERIALS AND METHODS

### Plant material and tissue collection

*Catharanthus roseus* (L.) cv. Dhawal (National Gene Bank, CSIR-CIMAP, Lucknow, India) plants were grown in a growth room maintained at 22–25°C with 16 h light and 8 h dark. MeJA treatment was performed as described in our previous study (Kumar et al., 2020). Briefly, the excised first pair leaves were dipped in 200 µM MeJA solution or water (control) containing the same amount of DMSO without MeJA. Samples were collected at different time points and stored at 80°C. For tissue-specific expression analysis, roots, stems, leaves, siliques and flowers were collected separately from 8-week-old plants, frozen in liquid nitrogen and stored at 80°C until further analysis. For VIGS/transient over-expression experiments, seeds were germinated and grown as mentioned above. Four-six leaf staged seedling were used for VIGS experiment and plants of 6-8 week old having 8-10 leaves were used for transient overexpression.

### Mining of CrROPs from MPGR database

To identify potential genes encoding ROPs in *C. roseus,* a search was carried out in the MPGR (http://medicinalplantgenomics.msu.edu) database initially by using ROPs and RACs of other plants as query and then later by blast search against *C. roseus* genome sequence (Kellner et al., 2015). Forty annotated ROP sequences were obtained and among them the sequences for which FPKM value was not assigned were excluded. After eliminating duplicate and pseudo sequences, a total of 6 ROP GTPase genes were finalized from the *C. roseus* genome.

### Phylogenetic analysis

For phylogenetic analysis, amino acid sequences of characterized Rac/ROPs from different plant species were retrieved from the National Center for Biotechnology Information (NCBI) data base (www.ncbi.nlm.nih.gov). The deduced amino acid sequences of CrROPs were compared with other Rac/Rop proteins obtained from the NCBI database. The phylogentic tree was constructed by neighbor-joining method using MEGA11 software(Tamura et al., 2007). Multiple sequence alignment of CrROP proteins was performed using Clustal (Thompson et al., 1994) with default parameters through EMBnet (http://www.ch.embnet.org/software/ClustalW.html).

### Protein expression, purification, and GTPase assay

For expression of recombinant CrROP proteins, the open reading frame of each gene was amplified from leaf cDNA using gene specific forward and reverse primers having suitable restriction sites (Table S3). The amplicons were cloned into pJET1.2/vector and the sequences were verified by Sanger sequencing. The confirmed *CrROPs* were restriction digested from pJET1.2-derived constructs using specific restriction enzymes and subcloned into pET28a. The resulting pET28a::CrROP constructs were transformed into *E. coli* Rosetta-2 competent cells and used for protein expression. Induction, harvesting and protein purification were performed as described earlier (Rai & Nagegowda, 2021) (Dwivedi et al., 2022) with slight modification. Briefly a single colony was inoculated into 10 mL Luria broth supplemented with kanamycin and chloramphenicol and inoculated overnight at 37 °C. From the overnight grown culture 5 mL was taken and added to 500 mL Luria broth containing the appropriate antibiotics and grown at 37°C till the absorbance at OD_600_ was 0.4. For protein induction, 50 µL of 1 M isopropyl-1-thio-β-D-galactopyranoside (IPTG) was added to the culture to get a final concentration of 1 mM and inoculated at 18 °C for 16 h. Protein purification by affinity chromatography on Ni-NTA agarose was performed according to manufacturer’s instruction. Protein concentration was determined following Bradford method (Bradford, 1976). GTPase assay was performed by using ATPase/GTPase Activity Assay Kit (Sigma-Aldrich) as per the manufacturer’s instructions.

### Protein prenylation assay

In vitro prenylation assays were performed as described in Dutilleul et al. (2016). Briefly, recombinant purified 6His-tagged CrROPs were incubated with C. roseus recombinant PFT or PGGT-I (Courdavault et al., 2005) with the corresponding NBD(NitroBenzoxadiazol)- modified fluorescent prenyl substrates (respectively NBD-FPP or NBD-GGPP, Jena Biosciences). GSTs equipped with a farnesylatable CAIM (GST-CAIM) motif or geranylgeranylatable CIIL motif (GST-CIIL) (Courdavault et al., 2005) were used as positive controls. In vitro prenylation reactions were composed of Tris 50 mM pH 7.5, ZnCl2 10 μM, MgCl2 5 mM, DTT 0.5 mM, NaCl 50 mM, NBD-FPP/GGPP 4 μM, 2 μg of protein prenyltransferase and 0.5-1 μg of recombinant CrROPS. After 1 h at 30°C, the reaction was stopped by the addition of Laemmli buffer and 5 min incubation at 95°C. Proteins were separated by SDS-PAGE and fluorescent proteins were visualized using the ChemiDocTM MP Imaging system (Bio-Rad). The protein gel was then stained in Coomassie blue solution.

### Analysis protein-protein interactions using yeast two-hybrid assay

For yeast two hybrid assay, the full-length sequence of *CrROP* genes (*CrROP2, CrROP3* and *CrROP5*) was cloned in-frame with the activation domain of pGADT7 (prey) vector, whereas *CrPGGT-I* was cloned in-frame with the DNA binding domain of pGBKT7 (bait) vector. The resulting constructs were transformed into mating yeast strains AH109 and Y187. Accordingly, AH109 harboring pGADT7::*CrROP2/3/5* and Y187 harboring pGBKT7::*CrPGGT-I* were selected on SD/-Leu and SD/-Trp media, respectively. Single colony from each of the above plates was co-inoculted into 1 ml of YPD medium and grown overnight at 30°C. About 100 ul of overnight grown culture was plated on SD/-Leu/-Trp medium and incubated at 30°C for two days. For determining the protein-protein interaction, the diploid yeast colonies grown on SD/-Leu/-Trp medium were further streaked on SD/- Leu/-Trp/-His/-Ade medium. For negative control the bait vector (pGBKT7::*CrPGGY-I*) and prey vector (pGADT7::*CrROPs*) were mated with the reporter strain containing the empty AD and DNA-BD vectors, respectively.

### Generation of silencing and overexpression constructs

The pTRV1 and pTRV2 vectors used for generating VIGS constructs were procured from The Arabidopsis Information Resource (TAIR), USA. The 500 bp fragments of *CrROP2/3/5* were amplified from leaf cDNA by PCR using gene specific primers having *Eco*RI restriction site at 5′ end of each primer (Table S3). The resulting PCR product was purified and sub- cloned into pJET1.2/vector and sequences were confirmed by nucleotide sequencing. The confirmed fragment was then cloned into pTRV2 vector digested with *EcoR*I. The resulting pTRV2::*CrROP*s plasmid was confirmed by digestion. For generation of overexpression constructs, the open reading frame of *CrROP2/3/5* were PCR amplified using leaf cDNA with specific forward and reverse primers (Table S3). A truncated *CrROP3* and *CrROP5* constructs lacking geranylgeranylation motif was generated by deleting 12 nucleotides from the 3’ end corresponding to CSIL motif. The amplicons were cloned into pJET1.2/ cloning vector for sequence confirmation, and then sub-cloned into *Xba*I and *Sac*I sites of pBI121 binary vector under the control of the 35S promoter of Cauliflower mosaic virus (CaMV) to form pBI121::*CrROP2/3/5* and pBI121:Δ*CrROP3/5* constructs (Figure S6& Figure 8). The VIGS and overexpression constructs were then mobilized into *Agrobacterium tumefaciens* GV3101 competent cells by freeze thaw method.

### VIGS and transient overexpression of *CrROPs*

VIGS and transient overexpression was performed according to Kumar *et. al.* (2015; 2020). Briefly the overnight grown *Agrobacteria* cultures harboring pTRV1 and pTRV2::*CrROP2/3/5* or pTRV2::*CrPDS* or pTRV2 (EV) constructs were resuspended in MES buffer (MES, MgCl_2_ and Acetosyringone) and mixed in 1:1 ratio before infiltration. Plants were infected by pricking below the apical meristem using a dissecting needle and kept in dark for 48 h. Plants were shifted to a growth chamber (22 °C, 75% humidity, 16-8h light cycle). Leaves from pTRV2::*CrROP2/3/5* were harvested 3-4 weeks post infiltration, when albino phenotype developed in the first 2 pairs of leaves in *CrPDS* infected plants, and stored in -80°C for further use. Transient overexpression was carried out following Kumar *et. al.* (2020) using *Agrobacteria* harboring pBI121::*CrROP2/3/5*, pBI121::Δ*CrROP3/5* and pBI121 (control) constructs. *Agrobacteria* with p19 (suppressor of RNA silencing) construct was mixed with pBI121::*CrROP2/3/5* or pBI121::Δ*CrROP3/5* or pBI121 in 1:1 ratio prior to infiltration.

### Generation of *C. roseus* transgenic plants overexpressing *CrROP2* and *CrROP5*

*C. roseus* transgenic plants were generated by tissue culture independent *Agrobacteium-* mediated transformation protocol as described earlier(S. R. Kumar et al., 2020). Agro- inoculated plants were PCR-screened using 35S forward and CrROP-specific reverse primers. Additionally, primers specific to the *Agrobacterium* chromosomal virulence gene (ChvA) were used to detect potential agro-contamination. Plants that exhibited no amplification or showed the presence of Agrobacterium contamination bands were excluded from further analysis. Plants that produced specific band only in 35S forward and CrROP-specific reverse primers combination were used for further analysis. T1 plants were obtained by PCR-screening the seedlings generated from seeds of T0 plants (Figure S7).

### Gene expression measurements

For tissue specific expression analysis, total RNA from leaves, roots, stems, siliques, flower buds and flower was extracted using Trizol reagent (Sigma Aldrich, USA) following the manufacturer’s instructions. Similarly, for determining the expression of genes in MeJA elicitation, VIGS, and overexpression experiments, leaves were collected and used for total RNA extraction. cDNA was synthesized using High-capacity cDNA Reverse transcription kit (Applied biosciences) following manufacturer’s instruction, and quantitative reverse transcription-polymerase chain reaction (qRT-PCR) was carried out as described previously (Rai et al., 2013; Kumar *et. al.,* 2020). The primers used for quantification of *CrROP* transcripts were designed outside of the gene region cloned in pTRV2. *CrN227* was used as endogenous control. The PCR conditions were as follows: 94°C for 10 min for one cycle, followed by 40 cycles of 94°C for 15 sec, and 60°C for 1 min. Fold-change differences in gene expression were analysed using the comparative cycle threshold (Ct) method. Relative quantification was carried out by calculating Ct to determine the fold difference in gene expression [ΔCt target – ΔCt calibrator]. The relative level was determined as 2_ΔΔCT.

### Metabolite analyses and quantification

Secologanin and MIAs were extracted as described earlier in Kumar *et. al.,* 2018. Additionally phenylpropanoid was extracted and analysed according to Pandey *et. al.,* 2014 with slight modification. HPLC (Model: SCL-10AVP, Shimadzu, Japan) equipped with C18 symmetry reverse-phase column (5 μM, 250 9 4.6 mM; Waters, Milford, MA, USA) was used for analysis. Data were recorded at 238 nM for secologanin, 254 nM for indole alkaloids. The samples were quantified in reference to the peak areas obtained from authentic standards (Sigma-Aldrich), and were expressed as relative content (%).

### Analysis of subcellular localization of CrROPs

The subcellular localization of CrROPs was studied by creating a GFP fusion protein using the p326sGFP vector. The ORF of each CrROP was amplified by PCR using leaf cDNA with gene-specific forward and reverse primers consisting of *Bsr*GI/*Not*I sites (Table S1). The resulting PCR product was cloned into pJET1.2 vector for sequence confirmation followed by sub-cloning into the *Bsr*GI/*Not*I sites of p326–sGFP vector yielding the CrROP-sGFP construct. Plasmids p326::CrROP-sGFP, PM-mCH, nuc-mCH and p326::sGFP were used for transient transformations of *C. roseus* cells by particle bombardment and fluorescence imaging were performed following the procedures previously described (Foureau et al., 2016).

### Statistical analysis

Average means, standard error (SE) and number of replicates (biological and technical) for each experiment was used to determine the statistical significance using GraphPad9 Prism software. The statistical significance of differences between control and treated samples were tested using one-way and two-way ANOVA with Sidak test.

## ACKNOWLEDGMENTS

This work was partly supported by CSIR FBR project MLP0003 to D.A.N. D.P.B., and A.S. are the recipients of a research fellowship from University Grants Commission (UGC). D.P. is the recipient of a research fellowship from Department of Biotechnology (DBT). D.P.B and D.A.N. thanks Dr. Ajit K. Shasany for hosting D.P.B. in his lab for short duration during the AcSIR course work at CSIR-CIMAP Lucknow. The authors are thankful to Dr. Ramu Vemana for sparing pDEST22 and pDEST32 vectors. The authors also express their sincere gratitude to the Director, CSIR-CIMAP for support throughout the study. Institutional communication number for this article is CIMAP/PUB/2023/131. Authors declare no conflict of interest.

## AUTHOR CONTRIBUTIONS

D.P.B., A.S., P.L.C., I.C., L.B., A.K., and D.P., performed the experiments. D.P.B., A.S., P.L.C., I.C., L.B., A.K., A.R., S.C.V., E.D., V.C., and D.A.N. analyzed the data. D.A.N. conceived and coordinated the research. D.P.B., A.S., A.R., and D.A.N. wrote the manuscript.

