## Supplemental Information for "Rho of plant GTPases with geranylgeranylation motif modulate monoterpene indole alkaloid biosynthesis in *Catharanthus roseus*"

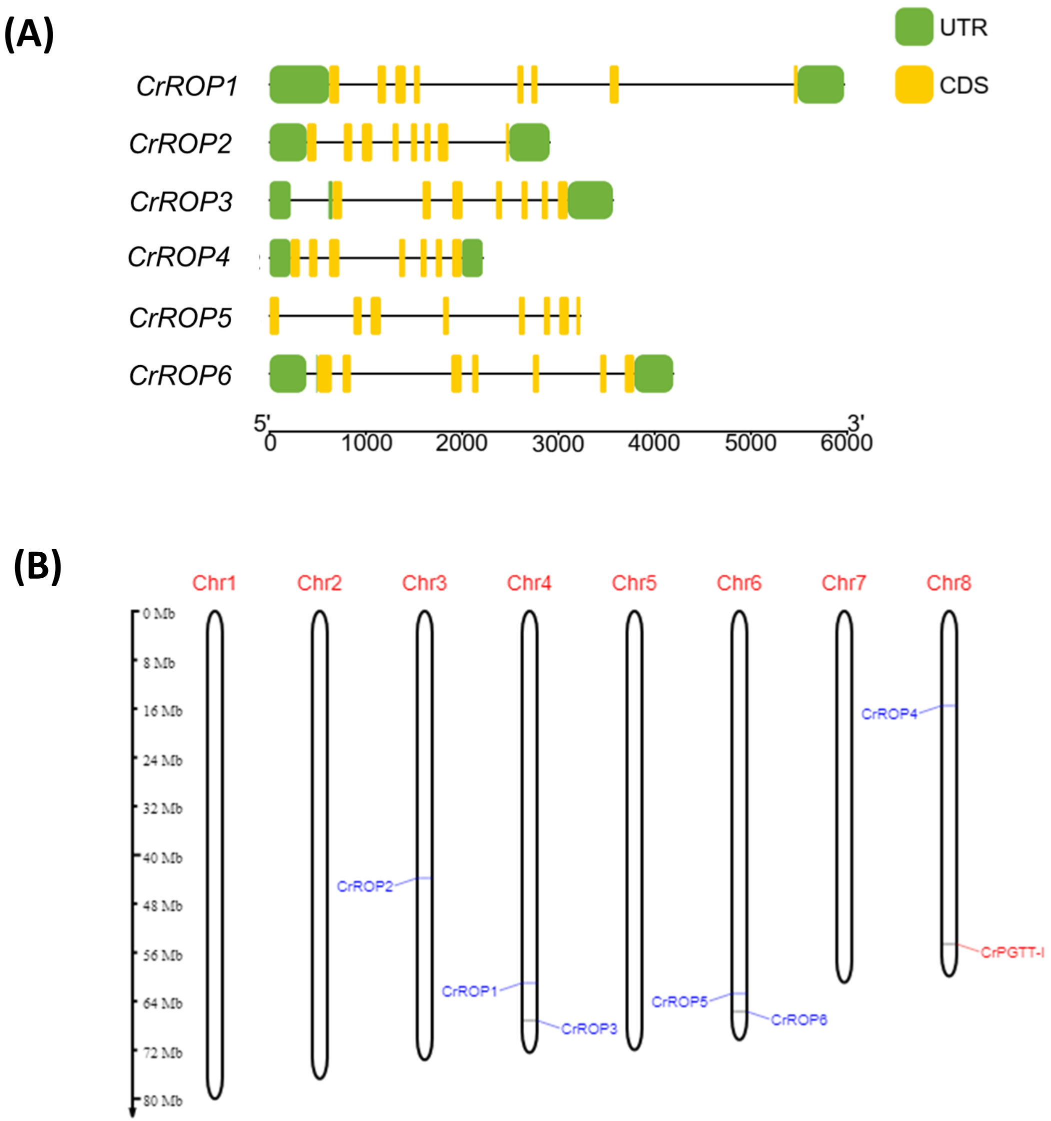


**Figure S1** (A) Schematic gene structure of *CrROPs* depicting exons (in yellow) and introns (black line) along with untranslated region (UTR, in green). The gene structure was made using TB tools (Chen et al., 2020). (B) Diagram showing the location of *CrROPs* and *CrPGTT-I* in *Catharanthus roseus* genome as obtained from MapGene2Chrom v2 (http://mg2c.iask.in/mg2c_v2.0/).

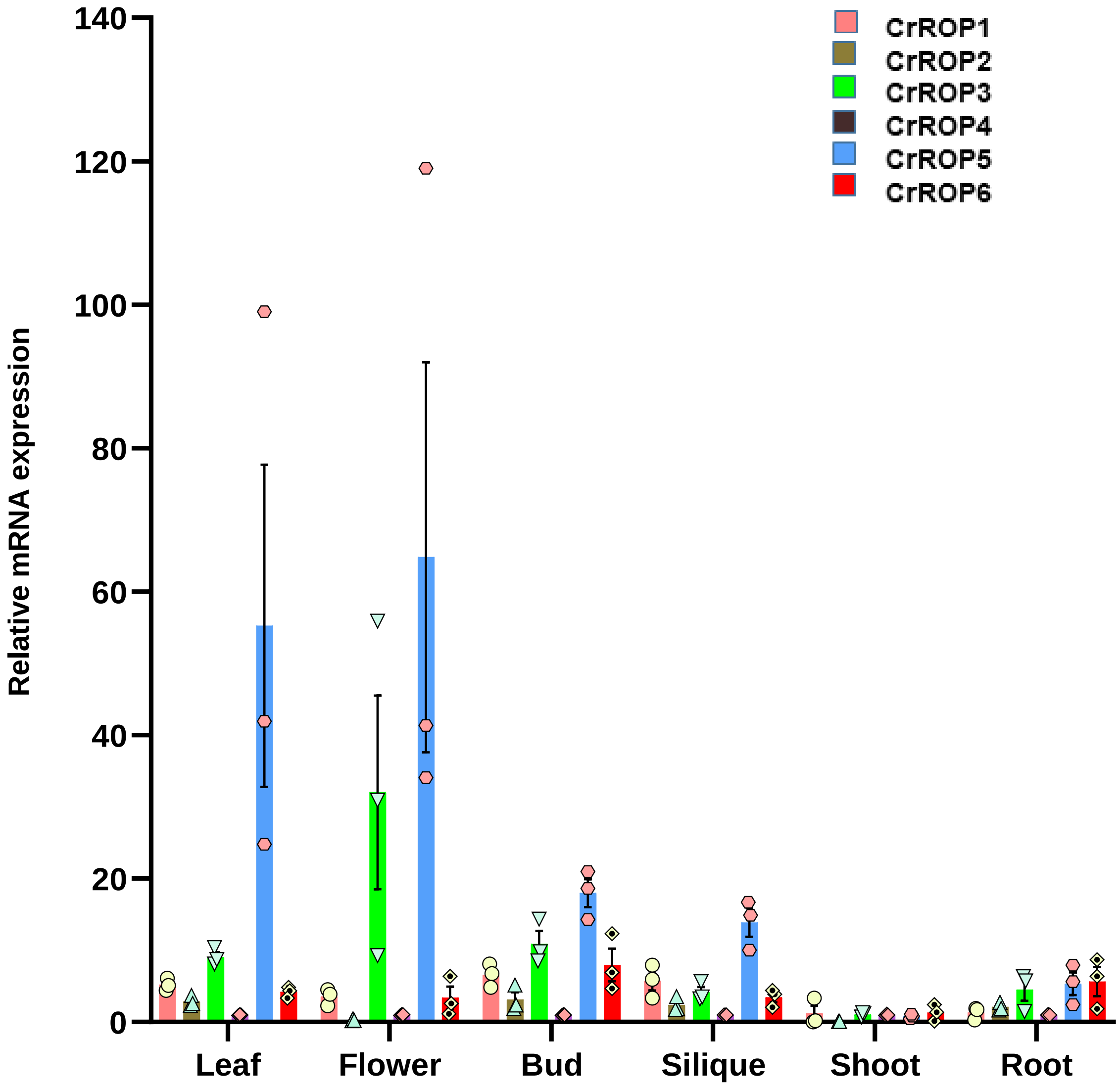


**Figure S2**. Relative transcript abundance of *CrROPs* in different tissues of *C. roseus. The e*xpression of CrROPs was measured in relation to the least expressor CrROP4 which was set to one.

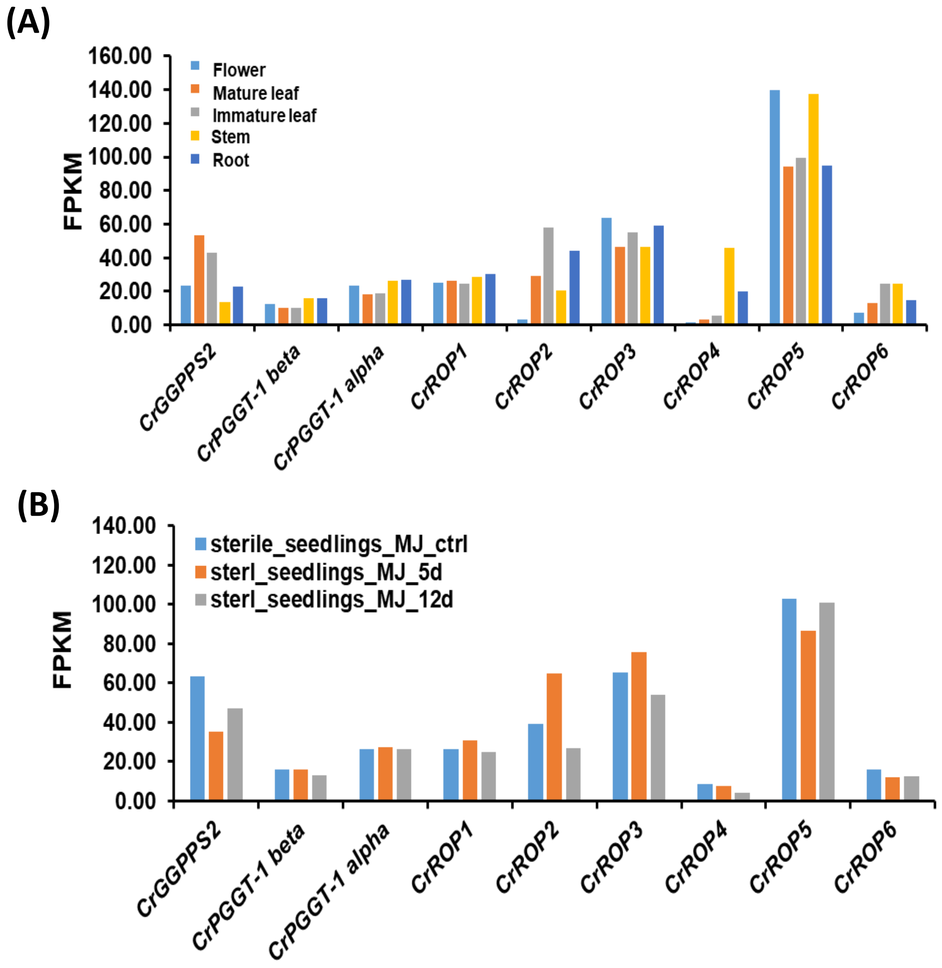


**Figure S3.** *In silico* gene expression of *CrROPs*, *CrPGGT* and *CrGGPPS2* in different tissues (A), and in response to MJ (MeJA) in seedlings (B). The FPKM values were obtained from MPGR database (http://mpgr.uga.edu/)

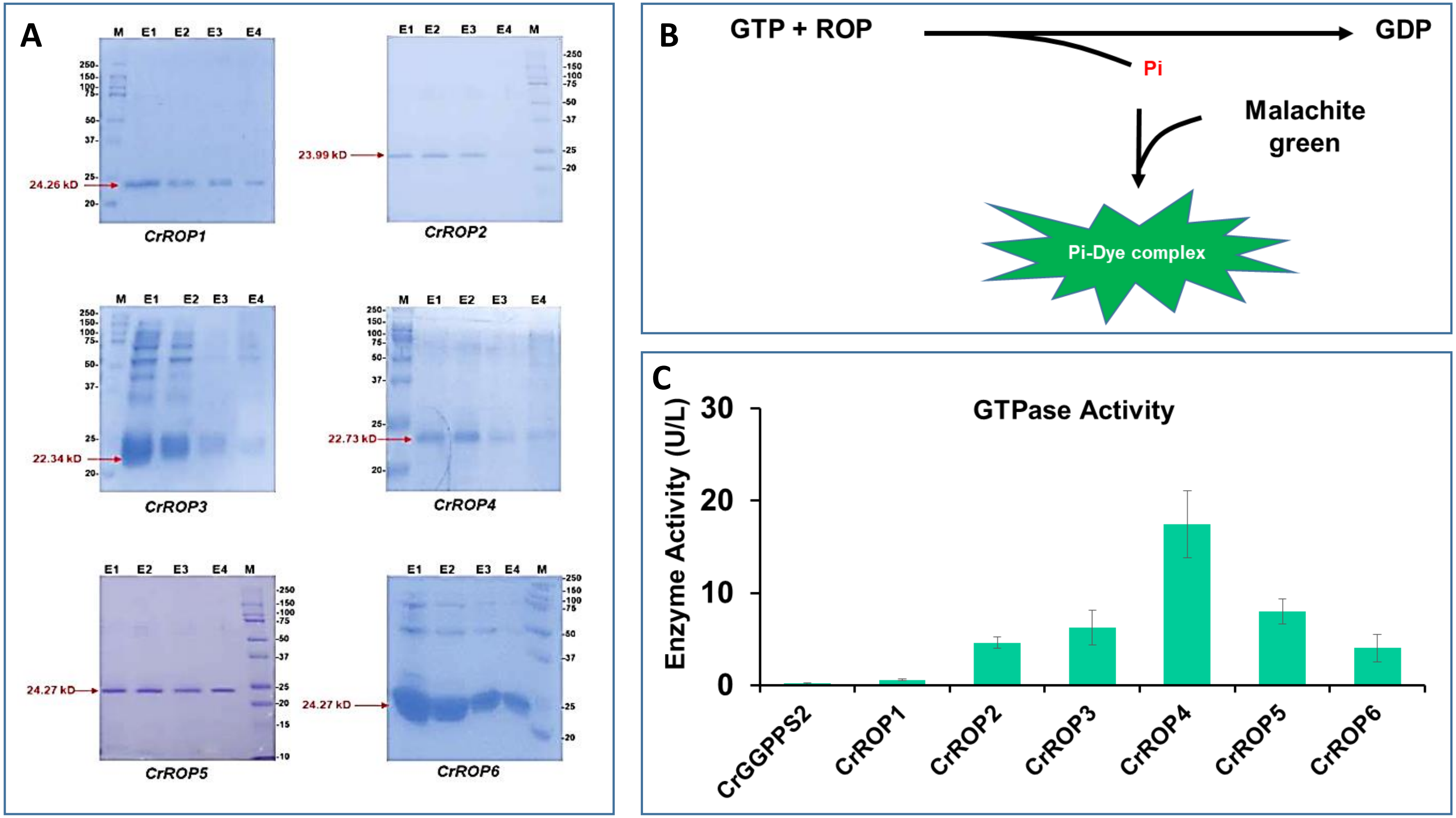


**Figure S4.** (A) SDS-PAGE analysis of partially purified recombinant CrROPs expressed in *E. coli* that were used for GTPase assay. Abbreviation; E1, elute 1; E2, elute 2; E3, elute 3; E4, elute 4; M, precision plus proteinTM dual colour marker. (B) Schematic GTPase reaction in presence of GTP and CrROP. (C) GTPase assay using Ni-NTA purified recombinant CrROPs. GTPase assay with GTP as a substrate was performed using purified recombinant CrROPs. The assay was done in three replicates and the enzyme activity is represented as U/L. Purified CrGGPPS2 was used a control.

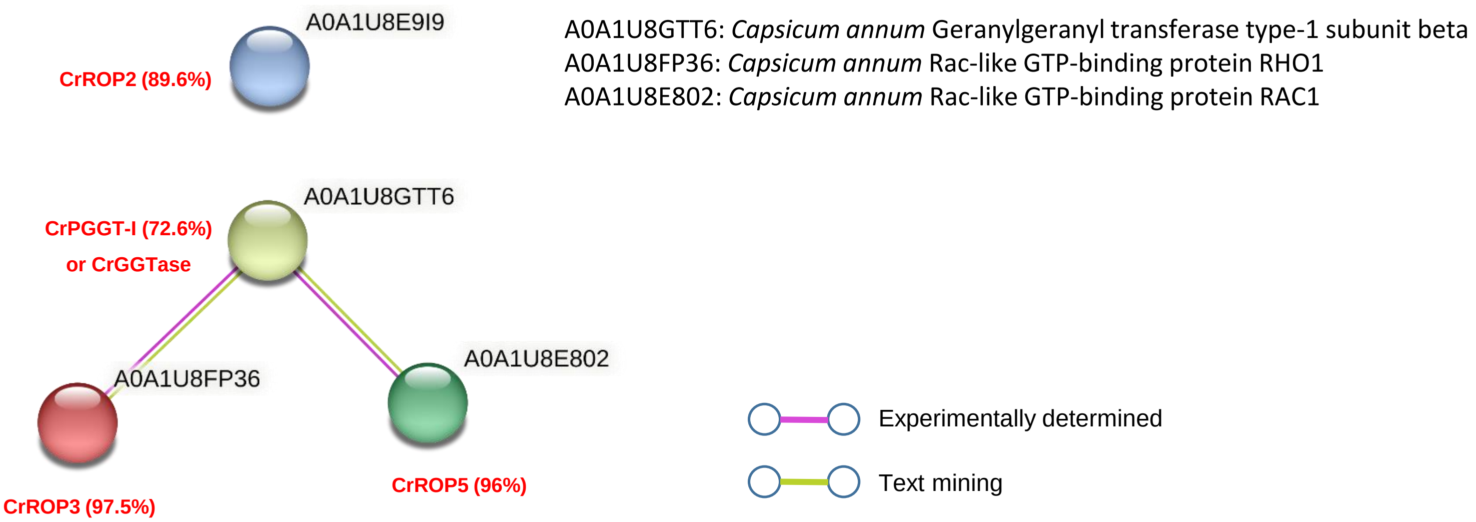


| node1 | node2 | node1 accession | node2 accession | node1 annotation | node2 annotation | score |
|---|---|---|---|---|---|---|
| CrGGTase | CrROP3 | A0A1U8GTT6 | A0A1U8FP36 | *Geranylgeranyl transferase type-1 subuni...* | *Rac-like GTP-binding protein RHO1* | 0.871 |
| CrGGTase | CrROP5 | A0A1U8GTT6 | A0A1U8E802 | *Geranylgeranyl transferase type-1 subuni...* | *Rac-like GTP-binding protein RAC1* | 0.871 |
| CrROP3 | CrGGTase | A0A1U8FP36 | A0A1U8GTT6 | *Rac-like GTP-binding protein RHO1* | *Geranylgeranyl transferase type-1 subuni...* | 0.871 |
| CrROP5 | CrGGTase | A0A1U8E802 | A0A1U8GTT6 | *Rac-like GTP-binding protein RAC1* | *Geranylgeranyl transferase type-1 subuni...* | 0.871 |

**Figure S5.** *In silico* analysis of protein-protein interaction between CrPGGT-I and CrROP2/3/5 using online prediction software STRING 11.5. tool (https://string-db.org/). (A) Green and purple lines indicate possible interaction of CrPGGT-I (homolog of *Capsicum annum* PGGT-I) with CrROP3 (homolog of *C. annum* RHO1) or CrROP5 (homolog of *C. annum* RAC1) . (B) Score for *in silico* interaction.

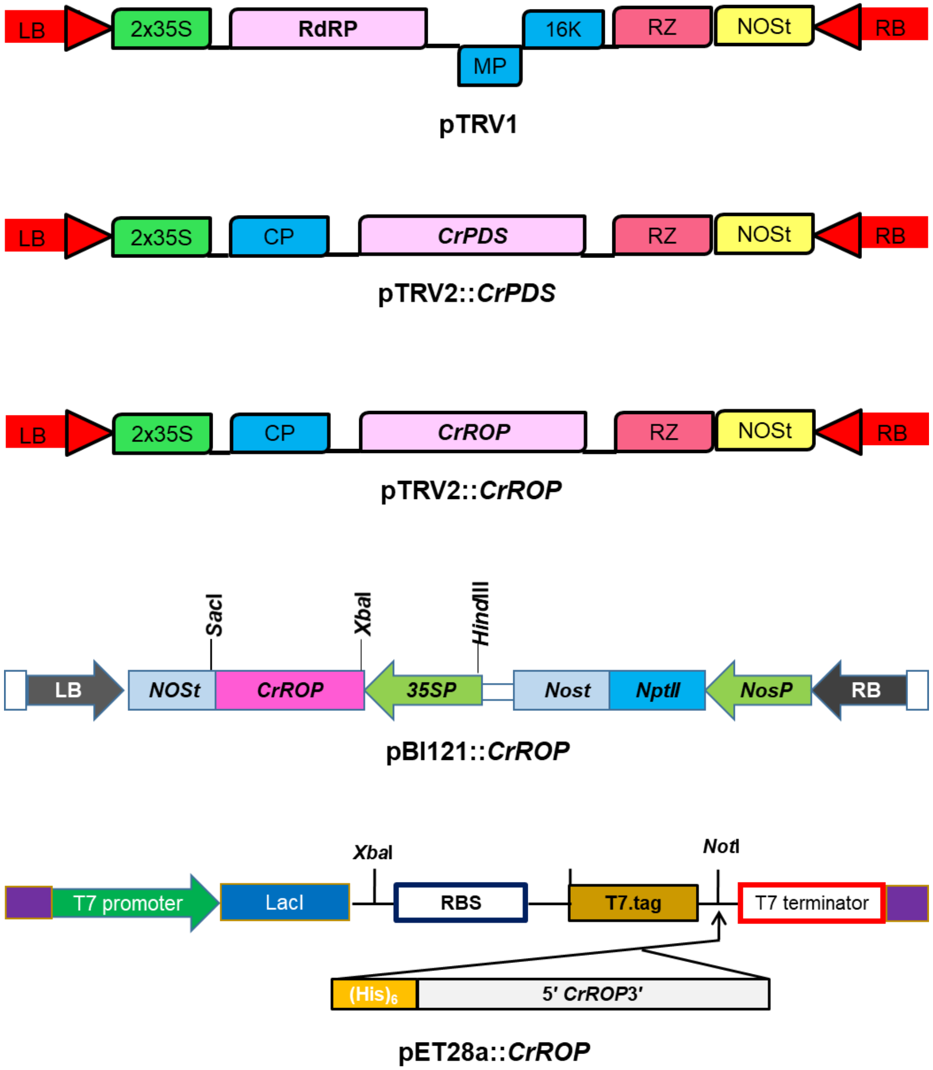


**Figure S6.** Vector maps of pTRV1, and pTRV2- pBI121- and pET28a-derived VIGS and overexpression constructs used in the study.

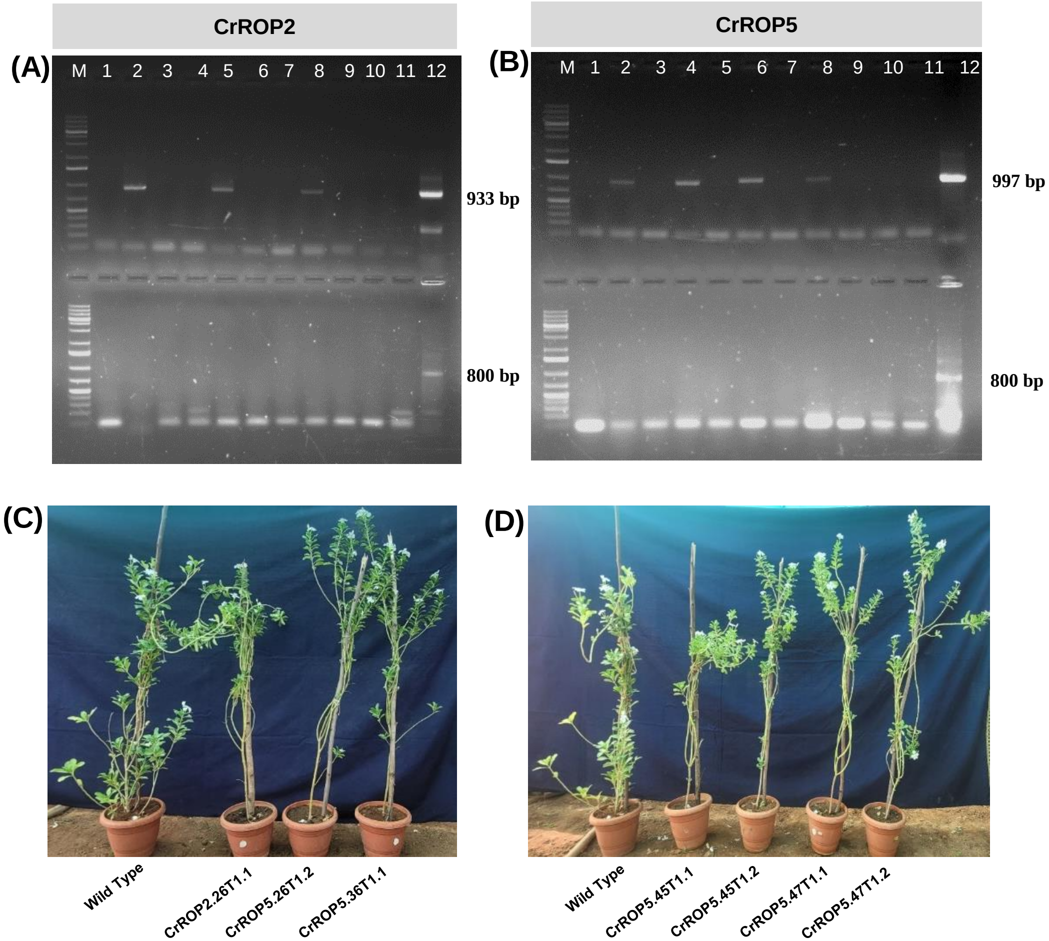


**Figure S7.** PCR-screening of plants generated from seeds of in planta transformed T0 *CrROP2* and *CrROP5* plants. (A) Top lanes: Agarose gel-electrophoresis showing amplification using CaMV35S-specific forward primer and *CrROP2*-specific reverse primer; (M): ladder; (2,5 & 8): single band corresponding to *CrROP2* confirming the transgenic nature of plants; (11): WT negative control; (12): Plasmid DNA pBI121::CrROP2 - positive control. Bottom lanes: Amplification using chv-F forward primer and chv-R reverse primer; (M): ladder; (1-10): absence of amplification at 800 base pairs confirms no Agro-contamination and transgenic nature of the plants; (11) WT negative control; (12): Agrobacterium GV3101 strain-positive control. (B) Top lanes: Amplification using CaMV35S-specific forward primer and CrROP5-specific reverse primer; (M): ladder; (2,4,6 & 8): single band corresponding to CrROP5 confirming the transgenic nature of plants; (11): WT negative control; (12): Plasmid DNA pBI121::CrROP2 - positive control. Bottom lanes: Amplification using chv-F forward primer and chv-R reverse primer; (M): ladder; (1-10): absence of amplification at 800 base pairs confirms no Agro-contamination and transgenic nature of the plants; (11) WT negative control; (12): Agrobacterium GV3101 strain-positive control. *In planta* transformed T1 *CrROP2* and *CrROP5,* and untransformed wild type *C. roseus* plants (C&D).

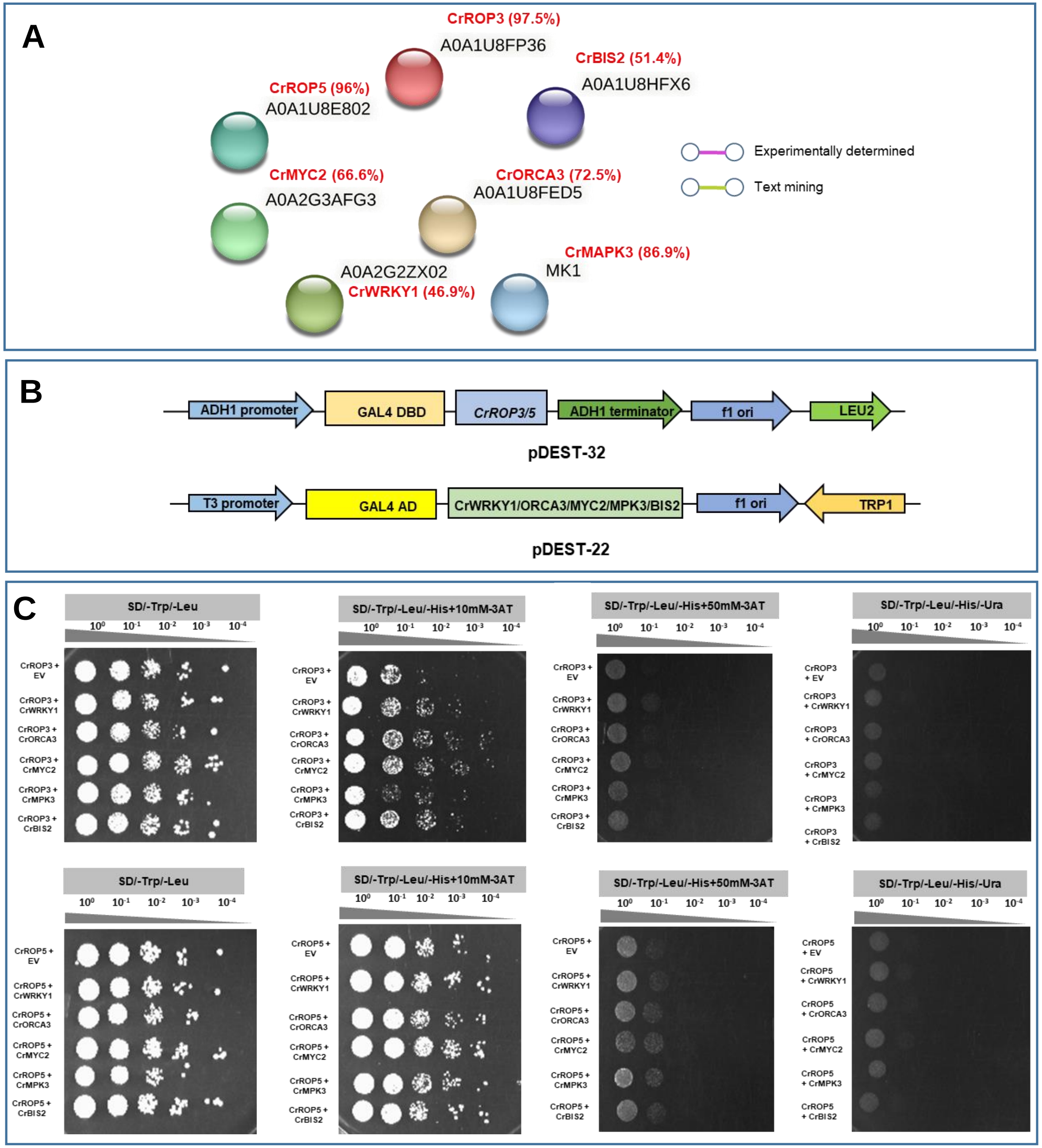


**Figure S8.** Yeast two-hybrid assay to determine the interaction between CrROP3 or CrROP5 and different transcription factors involved in regulation of MIA biosynthesis that were affected in CrROP3/CrROP5 VIGS and overexpression.

**Table S1.** Accession numbers and gene annotations of the sequences used in phylogenetic analysis.

| S. No. | Accession number | Gene Annotation | Plant Name |
|---|---|---|---|
| 1. | UHU71013.1 | CrROP1 | *Catharanthus roseus* |
| 2. | UHU71014.1 | CrROP2 | *Catharanthus roseus* |
| 3. | UHU71015.1 | CrROP3 | *Catharanthus roseus* |
| 4. | UHU71016.1 | CrROP4 | *Catharanthus roseus* |
| 5. | UHU71017.1 | CrROP5 | *Catharanthus roseus* |
| 6. | UHU71018.1 | CrROP6 | *Catharanthus roseus* |
| 7. | AEE78776.1 | AtRop1 | *Arabidopsis thaliana* |
| 8. | NP_173437.1 | AtRop2 | *Arabidopsis thaliana* |
| 9. | AEC06690.1 | AtRop3 | *Arabidopsis thaliana* |
| 10. | Q38937 | AtRop4 | *Arabidopsis thaliana* |
| 11. | AEE86595 | AtRop5 | *Arabidopsis thaliana* |
| 12. | Q38912 | AtRop6 | *Arabidopsis thaliana* |
| 13. | AED95321 | AtRop7 | *Arabidopsis thaliana* |
| 14. | AEC10456 | AtRop8 | *Arabidopsis thaliana* |
| 15. | NP_194624.1 | AtRop9 | *Arabidopsis thaliana* |
| 16. | NP_001327655.1 | AtRop10 | *Arabidopsis thaliana* |
| 17. | ANM69349.1 | AtRop11 | *Arabidopsis thaliana* |
| 18. | NP_001388456.1 | OsRac1 | *Oryza sativa* |
| 19. | NP_001407630.1 | OsRac2 | *Oryza sativa* |
| 20. | NP_001403809.1 | OsRac3 | *Oryza sativa* |
| 21. | NP_001389729.1 | OsRac4 | *Oryza sativa* |
| 22. | NP_001388946.1 | OsRac5 | *Oryza sativa* |
| 23. | NP_001105197.1 | OsRac6 | *Oryza sativa* |
| 24. | NP_001403453.1 | OsRac7 | *Oryza sativa* |
| 25. | NP_001104929.1 | ZmROP1 | *Zea mays* |
| 26. | NP_001105615.1 | ZmROP2 | *Zea mays* |
| 27. | AAD34357.1 | ZmROP3 | *Zea mays* |
| 28. | CAB96794.1 | ZmROP4 | *Zea mays* |
| 29. | NP_001105737.1 | ZmROP5 | *Zea mays* |
| 30. | CAB96793.1 | ZmROP6 | *Zea mays* |
| 31. | CAB96792.1 | ZmROP7 | *Zea mays* |
| 32. | NP_001105063.1 | ZmROP8 | *Zea mays* |
| 33. | NP_001105197.1 | ZmROP9 | *Zea mays* |
| 34. | AEQ62558.1 | Am-rac1 | *Aquilaria microcarpa* |
| 35. | AEQ62559.1 | Am-rac2 | *Aquilaria microcarpa* |
| 36. | ACI03398.1 | Sdrac1 | *Scoparia dulcis* |
| 37. | ACM07419.1 | Sdrac2 | *Scoparia dulcis* |
| 38. | BBN17866.1 | MpROP | *Marchantia polymorpha* |

**Table S1.** *In silico* subcellular localization prediction of CrROP2/3/5 using different software tools.

| Tools | CrROP2 | CrROP3 | CrROP5 |
|---|---|---|---|
| **Predotar** | None | None | None |
| **WoLF PSORT** | cytoplasm | chloroplast | chloroplast |
| **TargetP** | Any other location (other than chloroplast, mitochondria and secretory pathway) | Any other location (other than chloroplast, mitochondria and secretory pathway) | Any other location (other than chloroplast, mitochondria and secretory pathway) |
| **iPSORT** | Having a mitochondrial targeting peptide | Having a mitochondrial targeting peptide | Not having any of signal, mitochondrial targeting, or chloroplast transit peptides |
| **Plant-mPLoc** | Cell membrane. Cytoplasm. Nucleus | Cell membrane. Cytoplasm. Nucleus. | Cell membrane. Cytoplasm. Nucleus. |
| **TMHMM server** | No TMR | No TMR | No TMR |

**Table S3.** List of oligonucleotide primers used in this study.

| Name | Sequence (5´---------- 3´) |
|---|---|
| CrROP2-vigs-F | TCTAGACTGAAAGATGAGTGCTTC |
| CrROP2-vigs-R | CTCGAGGACATTCTGTTGTGTTTTGG |
| CrROP3-vigs-F | TCTAGAGAAGGAGTTGAGAAGTTCGG |
| CrROP3-vigs-R | CTCGAGACGAGCATTACAATGAGAATTG |
| CrROP5-vigs-F | TCTAGAGAAATTTCAGGGCTTGCC |
| CrROP5-vigs-R | CTCGAGGCACTCAATGTAAGCAGC |
| CrROP1-RT-F | AACCCAACAGAATGTGAAAGCA |
| CrROP1-RT-R | TGTTTCTGAGGTGGCTTGATTACT |
| CrROP2-RT-F | AGTGGCAAGAAAGAAAAGGCATA |
| CrROP2-RT-R | AAGCAGCACAACCTCCACAAA |
| CrROP3-RT-F | ACTGCTGGACAGGAGGATTACAA |
| CrROP3-RT-R | AGAGAAAAGGCCAAAATGAAAACA |
| CrROP4-RT-F | GGATTATGTGCCGACGGTTT |
| CrROP4-RT-R | CCCAGAGGCCCAAATTCAC |
| CrROP5-RT-F | CAGCCACCAAAGCAAAAGAAG |
| CrROP5-RT-R | TTCATCATTTTTTCAGCACTCACA |
| CrROP6-RT-F | TTCTGCTGCCTCTGCTACACA |
| CrROP6-RT-R | TTTTGCCAACAGCTCCATCTC |
| CrROP2-pBI12-F | GGTCTAGAATGAGTGCTTCAAAATTC |
| CrROP2-pBI121-R | GGAGCTCTCAAGCAGCACAACCTCC |
| CrROP3-pBI121-F | GTCTAGAATGAGCGCGTCGAGGTTC |
| CrROP3-pBI121-R | GAGCTCTCACAATATGGAGCAGGC |
| CrROP5-pBI121-F | GTCTAGAATGAGCGCCTCCAGGTTC |
| CrROP5-pBI121-R | GGAGCTCTCACAATATACTGCAAGCCCTC |
| ΔCrROP3-pBI121-F | GTCTAGAATGAGCGCGTCGAGGTTC |
| ΔCrROP3-pBI121-R | GAGCTCTCAGGCCTTTTGAGCCTTTCCC |
| ΔCrROP5-pBI121-F | GTCTAGAATGAGCGCCTCCAGGTTC |
| ΔCrROP5-pBI121-R | GAGCTCTCAAGCCCTCTGACCCTTTC |
| pGADT7::CrROP2-F | GCATATGATGAGTGCTTCAAAATTC |
| pGADT7::CrROP2-R | GGAGCTCTCAAGCAGCACAACCTCC |
| pGADT7::CrROP3-F | GCATATGATGAGCGCCTCCAGGTTC |
| pGADT7::CrROP3-R | GGAGCTCTCACAATATACTGCAAGCCCTC |
| pGADT7::CrROP5-F | GCA TAT GAC CAC TGC AAG ATT TAT C |
| pGADT7::CrROP5-R | GAGCTCTCAAGGAAAATGCAAGATG |
| pGBKT7::CrPGGTI-F | GAATTCATGGCGGACGACGATTACGCATG |
| pGBKT7::CrPGGTI-R | GGATCCCTAGAGAAAAAATCCAGTAGCAGC |
| CrN227 RT-F | TCCTTACGCCGCATTATCAG |
| CrN227 RT-R | AGATGAGACAGTAACGCCTTG |
| CrPGGTI RT-F | GTTGCATCTTTGCGCCTGAT |
| CrPGGTI RT-R | AAGCTCCACTCGAGAAGCATAGG |
| CrORCA1 RT-F | CATTCAGAAACGCGATAGAAGCT |
| CrORCA1 RT-R | TGAAGCCCTATAGTTCGGAAGATT |
| CrORCA2 RT-F | GCGGAAGATGCGGCATTA |
| CrORCA2 RT-R | CCGGAGCATTAGCAGAACCA |
| CrORCA3 RT-F | TGATTCCAGCTCGGAATTGAC |
| CrORCA3 RT-R | GGCTACCGCGTTTCTGTATTTC |
| CrG8H RT-F | CTTTCCAAAAAACACGGTCCAA |
| CrG8H RT-R | TCGCCATTGTTGATGAAGATATG |
| CrSLS RT-F | GAAAGGCAATTGCTGCCACT |
| CrSLS RT-R | ACCATGCCCAATCCAACACT |
| CrAS RT-F | GCGAACATTTGCAGATCCAT |
| CrAS RT-R | GGCCGATTTGTTATTGTTCC |
| CrSTR RT-F | TGCCACACAACTAGCCACAA |
| CrSTR RT-R | TCATGATTTCTTCCACACCTTCG |
| CrD4H RT-F | TGGCCTCAGTAGCAATTCAG |
| CrD4H RT-R | TCCATATTTCTCACTCGCTTCTC |
| CrT16H RT-F | ACTTCCAAGAGAATGTCGAGAAC |
| CrT16H RT-R | TCTATCCGGGTTAAATTTCTCAGG |

| Name | Sequence (5´---------- 3´) |
|---|---|
| CrMYC2 RT-F | TTGATTGCTGGGCCACAAG |
| CrMYC2 RT-R | TCGGTATCGGTCACCTCTTCA |
| CrSGD RT-F | CCATCGTTCATCGTCGAGATT |
| CrSGD RT-R | CACCCTCACACTGATAAGCAGATC |
| CrTDC RT-F | AGCCACTTGAAGCTGAGGAATT |
| CrTDC RT-R | CTTCGCTAAGGACCGGATATGTT |
| CrGES RT-F | ATTGATGGAAGCCCAAGAATTC |
| CrGES RT-R | ACCTTCTTAGAAAGAGATGGTGCC |
| CrWRKY RT-F | CGTACTTGGTCCCGACGATATT |
| CrWRKY RT-R | CGTCGGTGATTTCGGAACTAG |
| CrBIS1 RT-F | ATGGAATCAGTGGTGCTAGTGA |
| CrBIS1 RT-R | TTCAATTTCAGGGAGCTGTGAC |
| CrBIS2 RT-F | CCATTATTGCTGAGATGGAGAA |
| CrBIS2 RT-R | CTTCATTCAGTCTGCCATTGGT |
| CrMPK3 RT-F | AAGGGCAAAGACCTGATTGG |
| CrMPK3 RT-R | TCAGGCAAACTTGGTGGTTC |
| CrORCA4 RT-F | ATAGTAGTACTGCCGCCGAAAG |
| CrORCA4 RT-R | ATCTCCGCCGCAAATTTTCC |
| CrORCA5 RT-F | TCTTCCAACGGAGGTTAACGG |
| CrORCA5 RT-R | AATGTTGTCTCCAGGGCTTG |
| CrCPR RT-F | TAAGAACCGCCCTCACTCGTT |
| CrCPR RT-R | ATCGGCCTCATTTGGATCAG |
| CrDXR RT-F | CTCACTATCTTTTTGGCGCTGAA |
| CrDXR RT-R | CAAGACCGATGAATCCTGTGTCT |
| CrDXS RT-F | TTGATGGCCACAACATTGATG |
| CrDXS RT-R | TGCCTTTCTCGGTGACAACA |
| CrHDS RT-F | CCAACGCAGGACAGGTGAT |
| CrHDS RT-R | GGGAAACGGACATGAGAACAG |
| CrMECS RT-F | GCTGCCGTTACTCCTTCTAAGATT |
| CrMECS RT-R | CCAGGTTCCAAACGATGAAGA |
| CrROP1-pET-28a-F | GCATATGATGGCTGCAAATGCATCAAG |
| CrROP1-pET-28a-R | GGAGCTCTCACTTCAGACATACTAGTTTTC |
| CrROP2-pET-28a-F | GCATATGATGAGTGCTTCAAAATTC |
| CrROP2-pET-28a-R | GGAGCTCTCAAGCAGCACAACCTCC |
| CrROP3-pET-28a-F | GCATATGATGAGCGCCTCCAGGTTC |
| CrROP3-pET-28a-R | GGAGCTCTCACAATATACTGCAAGCCCTC |
| CrROP4-pET-28a-F | GCATATGATGAGCGCGTCGAGGTTC |
| CrROP4-pET-28a-R | GAGCTCTCACAATATGGAGCAGGC |
| CrROP5-pET-28a-F | GCATATGACCACTGCAAGATTTATC |
| CrROP5-pET-28a-R | GAGCTCTCAAGGAAAATGCAAGATG |
| CrROP6-pET-28a-F | GCATATGATGACTAGTACAAGTTCTGTTGC |
| CrROP6-pET-28a-R | GGAGCTCCTAGAGCAGAATACATGCTTT |
| CrWRKY1 GW F | GGGGACAAGTTTGTACAAAAAAGCAGGCTTAATGAACACCACGATGGCTTGG |
| CrWRKY1 GW R | GGGGACCACTTTGTACAAGAAAGCTGGGTTCAAAAGAGTAGAATGTGAAGGCAATGATTTGC |
| CrBIS2 GW F | GGGGACAAGTTTGTACAAAAAAGCAGGCTTAATGATGACGATGATGATGGATAATTCAGCA |
| CrBIS2 GW R | GGGGACCACTTTGTACAAGAAAGCTGGGTTGTCTGCCATTGGTGGACTGTTAAGA |
| CrMYC2 GW F | GGGGACAAGTTTGTACAAAAAAGCAGGCTTAATGAACCTATGGGGTACGACGAC |
| CrMYC2 GW R | GGGGACCACTTTGTACAAGAAAGCTGGGTTTACCAAGAGCCTCATCGAGTTTCC |
| CrMPK3 GW F | GGGGACAAGTTTGTACAAAAAAGCAGGCTTAATGGTTGATGCAAATATGGCCG |
| CrMPK3 GW R | GGGGACCACTTTGTACAAGAAAGCTGGGTTTGCATATTCTGGATTTAGAGCCAAGG |
| CrORCA3 GW F | GGGGACAAGTTTGTACAAAAAAGCAGGCTTAATGTCCGAAGAAATCATTTCCGTCTCA |
| CrORCA3 GW R | GGGGACCACTTTGTACAAGAAAGCTGGGTTATATCGTCCCTTCTTCCTTCCTCC |
| CrROP3GWFP | GGGGACAAGTTTGTACAAAAAAGCAGGCTTAATGAGCGCGTCGAGGT |
| CrROP3GWRP | GGGGACCACTTTGTACAAGAAAGCTGGGTTCAATATGGAGCAGGCCTTTTGAGC |
| CrROP5GW FP | GGGGACAAGTTTGTACAAAAAAGCAGGCTTAATGAGCGCCTCCAGGT |
| CrROP5GWRP | GGGGACCACTTTGTACAAGAAAGCTGGGTTCAATATACTGCAAGCCCTCTGACC |
